# Changes in species composition of sessile communities on subtidal rock walls in the southern Gulf of Maine during four decades of warming

**DOI:** 10.64898/2026.03.01.708879

**Authors:** Breck A. McCollum, Jarrett E. K. Byrnes, Kenneth P. Sebens

## Abstract

Climate change is driving species range shifts and population change in density and location globally. Two theories behind these shifts, that species in the ocean are largely tracking climate velocities, and the concept of long-term temporal turnover, have garnered increased attention recently. However, research in marine ecosystems has largely focused on mobile species, namely commercially important fishes. Here we examine changes in sessile invertebrate and algal species on vertical surfaces, subtidal rock walls, in the southern Gulf of Maine (GOM), and to what extent these changes might have been driven by 42 years of warming. In part due to ocean circulation patterns in the GOM, the thermally-sensitive species in this community are unlikely to track climate velocities by moving laterally, and are therefore disappearing, moving into deeper water, or adapting to novel thermal conditions. We find that some species, including one of the previously competitive dominants, *Alcyonium siderium*, have become exceedingly rare at these sites. Two other competitive dominants, *Metridium senile* and *Aplidiiuam glabrum*, have also declined precipitously. Meanwhile, the blue mussel, *Mytilus edulis*, the non-native tunicate *Didemnum vexillum*, and a complex of erect bryozoans have become dominant space holders. Over the same period of time, average summer temperatures in the southern GOM increased by more than 3°C. Using occupancy derived thermal affinities, we find warm-affinity species increasing, while generally, cool and cold-affinity species are decreasing. All species which decreased in abundance normally occupy sites with temperatures below a mean of 17.4°C maximum summer temperatures. A few species did not change abundance despite the rapidly warming surface waters, indicating their broad tolerances and the importance of other biological processes in mediating community structure in the GOM. Overall, sessile rock wall communities in the southern GOM are transitioning to more thermally-tolerant species, most of which are not native to the Atlantic coast of North America.

## Introduction

Human-induced climate change has affected the global distribution of numerous terrestrial and aquatic species (Burrows et al., 2011; Lawlor et al., 2024; Lotze et al., 2022; McHenry et al., 2019; Sunday et al., 2012; Witman et al., 2023). Over the last 60 years, approximately 84% of all heat produced by global warming has been absorbed by the oceans (Barnett et al., 2005; Cheng et al., 2025), leading to range shifts in marine species as waters warm (Sunday et al., 2015). Shifts in water temperature have led to the temporal turnover (change in species composition) of communities (Pinsky et al., 2025) as the makeup of communities begin to tropicalize or deborealize (Chust et al., 2024). It has been predicted that North Atlantic species ranges will contract considerably and become patchier as ocean temperatures continue to warm (McHenry et al., 2019).

Range shifts include the expansion, contraction or lateral movement of a species or population distribution. The perimeter of a species’ range is often dynamic, and can shift based on several factors including temperature, salinity, competition, prey availability, and suitable habitat among others. Analogies have been made between range shift biology and invasion biology (Dunstan and Bax, 2007; Sorte et al., 2010), as species in both scenarios encounter novel environments and may have strong impacts on the existing communities. The distributions we observe today are mere ‘snapshots’ of species responding to past biological or climatological events (Dunstan and Bax, 2007). As climatological events such as temperature change increase, species able to migrate laterally along the coast in response will have an advantage under future environmental conditions. The ability of sessile species to shift their ranges depend on larval dispersal properties, direction of current flow, and the proximity of suitable habitat. When species are unable to move, they must adapt or face local extinction (Gaylord and Gaines, 2000; Burrows et al., 2011; Bates et al., 2014; Sunday et al., 2015; Witman et al., 2023).

Marine species are considered thermal-range conformers; i.e. their thermal tolerance closely matches their latitudinal range (Sunday et al., 2012), marine species, in particular, appear to track climate velocities (Pinsky et al., 2013) by lateral movement. Climate velocities are defined by Pinsky et al., 2013 as “the rate and direction that climate shifts across the landscape”. The rapid pace at which the climate has been changing in recent decades suggests that range shifts will be the dominant process affecting community structure and ecosystem functioning (Bates et al., 2014). A meta-analysis of 1700 species (both terrestrial and marine) found an average poleward range shift of 6.1 km/decade (Parmesan and Yohe, 2003). Similarly, Sorte et al., 2010 found that 75% of all marine species range shifts were in a poleward direction, consistent with climate change predictions of warming temperatures. Additionally, Pinsky et al., 2013 found that of 360 marine species, 74% shifted in the direction of their climate velocities. Therefore, as temperatures rise, species can be expected to migrate towards the poles (Barry et al., 1995; Holbrook et al., 1997; Lubchenco et al., 1993; Southward et al., 1995). Likewise, cold-affiliated species should decrease in abundance while warm-affiliated species should increase in abundance (Barry et al., 1995; Beaugrand et al., 2002; Holbrook et al., 1997; Southward et al., 1995).

The Gulf of Maine provides an ideal venue to study the effects of climatic warming on the ability of marine communities to respond to changing environmental conditions given its steep thermal gradient and rapidly warming waters. The GOM is a semi-enclosed, relatively shallow basin with an average water depth 150 m (Figure 1, inset). Much of the GOM seafloor is soft sediment, but the coastline is dotted with rocky outcrops and small islands. The GOM is fed by two major oceanic sources, arctic meltwater from the north, and warm Gulfstream waters from the south (Lotze et al., 2022; Record et al., 2023). In recent years, increased intrusion by Gulf Stream waters has caused sea surface temperature in the GOM to warm faster than 99% of the world’s oceans (Pershing et al., 2021, 2015). Additionally, circulation in the GOM generally flows in a southward, counter-clockwise direction (Lotze et al., 2022). This circulation pattern presents a large obstacle to range expansion by organisms living in the GOM.

**Figure 1.**
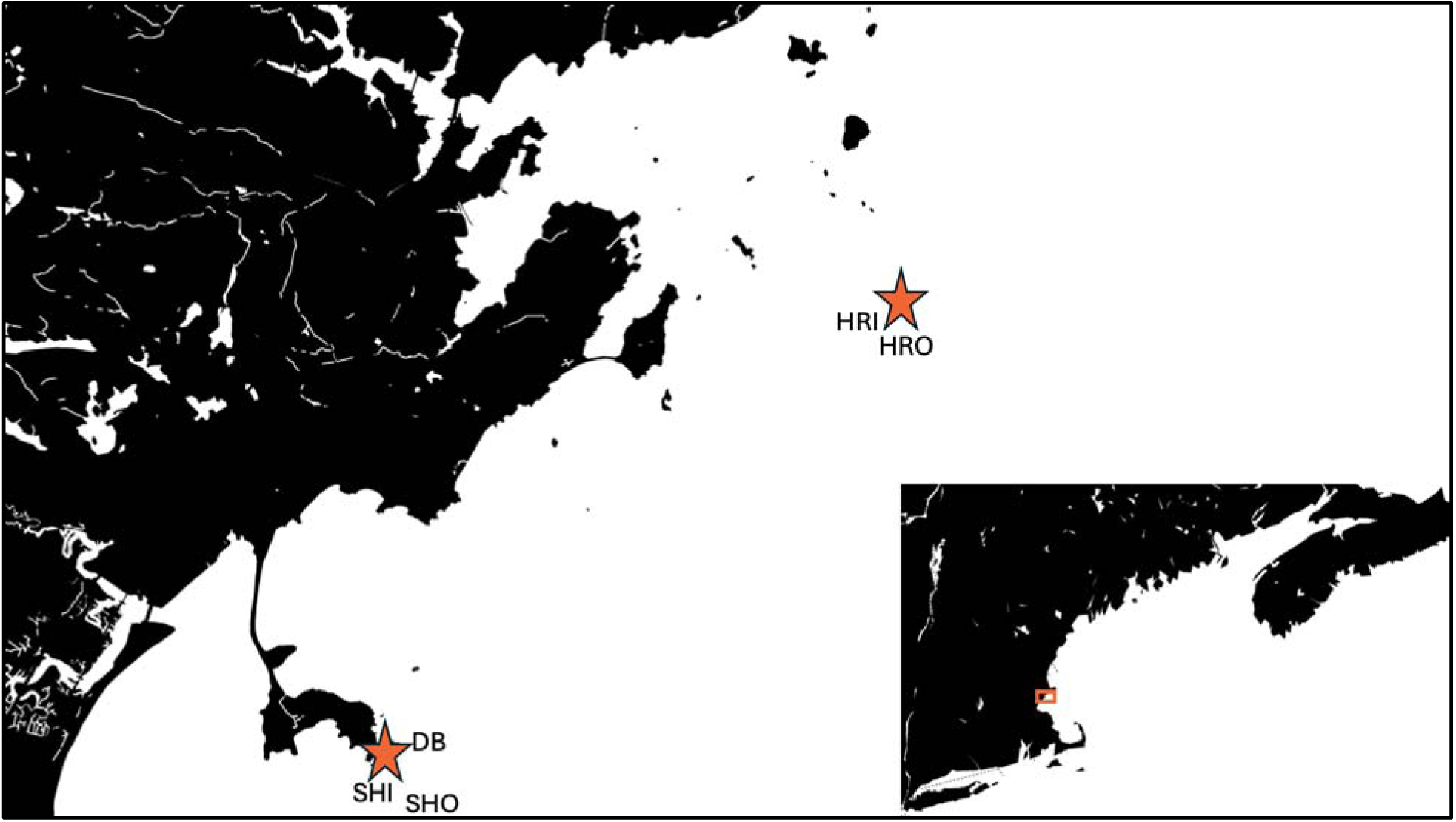
Map of study sites located within the Gulf of Maine (inset). Stars indicate the locations of Halfway Rock and Nahant, MA, each containing multiple study sites.

The shallow, rocky subtidal zone in the GOM can be summarized as algae-dominated horizontal benches or coralline algae and invertebrate-dominated vertical rock surfaces (walls) (Byrnes et al., 2024; Genovese and Witman, 1999; Leichter and Witman, 1997; Miller and Etter, 2011; Sebens, 1985, 1986; Steneck et al., 2013). The vertical surfaces of subtidal rock walls are generally light-limited beyond 3 m in depth and are therefore dominated by encrusting invertebrates rather than macroalgae (Miller and Etter, 2011; Sebens, 1985, 1986). Under low grazing pressure, these walls in the rocky subtidal zone reached a “climax” community state (1980s) with patches of the octocoral A*lcyonium siderium,* the colonial tunicate *Aplidium glabrum,* and the plumose sea anemone *Metridium senile,* growing over multiple species of crustose algae, encrusting bryozoans, sponges and other low-relief invertebrates (Sebens 1985, 1986). These species represent the hierarchical winners (i.e. competitive dominants) in terms of space occupation in an undisturbed environment, and each one can exclude the establishment of other sessile species and persist for long periods of time (Sebens 1985, 1986). The hierarchical structure of rock walls in the GOM includes very little bare rock; coralline algae and fleshy red crusts are often the settlement surfaces for most invertebrates in this community with a planktonic larval stage or crawling demersal larvae (e.g. *A. siderium*, Sebens, 1983). Intermediate levels in the sessile community competitive hierarchy include various sponges, tunicates, hydroids, and bryozoans, some of which now include very successful non-native species (Sebens 1985, 1986). Here we explore the response of this community of sessile invertebrates and algae to rapidly increasing temperature, in an environment with limited potential for range shifting, by documenting patterns of change over the last 42 years.

We utilized a novel data set for rock walls off Massachusetts, USA, initiated by one of us (Sebens) and assessed from 1978 to 2020 by SCUBA diving and quadrat photography, to examine how the sessile invertebrate community on subtidal rock walls has changed over time, and to examine whether these shifts are related to species responses to the background warming experienced by the GOM. We evaluated how the abundances of all component species, as percent cover on rock surfaces, have changed over time. We then related coefficients of change in abundance to derived species thermal preferences (i.e. their known temperature maxima). Broadly, we found that communities on rock walls in the rocky subtidal zone in the southern GOM are transitioning towards species assemblages that are more thermally-tolerant.

## Methods

### Survey History and Methodology

Data for these analyses comes from a long-term survey started in 1978 at two locations (Fig. 1) in the southern GOM - at Halfway Rock outside of Salem Sound, MA, USA (42.502573, -70.775086) and at East Point of Nahant, MA, USA (42.416867,- 70.906200). Multiple sites are present at both locations – Halfway Rock Inner (HRI), Halfway Rock Outer (HRO), Dive Beach (DB), Shag Rocks Inner (SHI), and Shag Rocks Outer (SHO). Sites were further divided into subsites (Supp. Material Table 1) spanning a depth gradient from 7-16 m. We took photographs of 4-6 25 x 35 cm marked permanent quadrats quarterly, covering each subsite, as detailed in Sebens 1985, 1986) and for this analysis we used use data from the summer (June - September). Quarterly sampling was maintained through 2005, with some omissions, and most later years had 1-3 samples per year in spring, summer or fall. We selected the summer season only, because that had the most consistent sampling over the 42-year period.

We analyzed photographic quadrats for percent cover of each organism to the lowest taxonomic level. From 1978 - 2007, we calculated percent cover using multiple acetate film sheets marked with 200 random points so subsequent photos of the same quadrat never had the same points used. We identified organisms to the lowest taxonomic level possible under each point. Some difficult to identify organisms were assigned to coarser taxonomic groups such as “Erect Bryozoan”. Points with a mat of amphipod tubes or a mass of mixed hydroids and bryozoans were assigned to “Tube Complex” or “Hydrozoan/Bryozoan Complex” classes respectively.

We reassigned points falling on the upper portion of a canopy forming organism or on mobile fauna to ensure we recorded data from primary substrate occupiers only. Canopy points were tracked separately for use in other analyses, as were counts of mobile species in each quadrat. Points falling on shadow, site markers, camera framer, rubble, shell hash or sediment were also excluded from the 200-point analysis. Starting in 2007, we processed images the same way using Coral Point Count (CPCe) (Kohler and Gill, 2006). In 2014, identifications were moved to CoralNet (Beijbom et al., 2015) which uses machine learning to identify species or categories under points for each image, but methods remained otherwise the same. These automated programs generate new unique sets of random points for every image. We processed point identification data from all three methods into percent cover for each photograph (https://github.com/jebyrnes/sebens_data_processing) dividing number of points per species or group by total number of identifiable and unobstructed substratum points assessed in an image (usually 100-200 points per photo).

### Temperature Change in the Southern GOM

We used loggers (Onset Hobo Tidbit data loggers) to record ocean temperature every 30 minutes at each site starting in 1996. To build a temperature time series for these sites, we selected the logger from each site, calculated monthly averages, and compared that data to the NOAA Global Ocean Data Assimilation Systems (GODAS) Potential temperature product at a depth of 15m (NOAA PSL, Boulder, Colorado, USA, https://psl.noaa.gov, Behringer 1998) using linear regression. We accessed the GODAS_15m data for a one grid cell (1/3° x 1°) centered on our sites. We then interpolated missing monthly logger values from 1980 to the present to construct a complete combined time series. From this time series, we calculated reference summer mean temperatures from the time periods of 1980-1985 and 2015-2020.

### Change in Species Abundances

We fit a model for each species or taxonomic group and examined its coefficient of change to determine the direction of change in percent cover of each species or taxonomic group over time, and its significance. We converted percent cover data to proportions and fit ordered beta regression models (Kubinec, 2023) using the glmmTMB package (Brooks et al., 2017) in R (Rstudio Team, 2016). Ordered beta regression is preferred for modeling proportion data that has an excess of zeros, ones, or both by estimating a cut point for each (Kubinec, 2023). This is necessary as the Beta distribution cannot accommodate zeros or ones, and zero-one inflated beta regression (ZOIB) requires a large number of extra parameters (Kubinec, 2023). For each species or taxonomic group, we modeled proportional cover as a function of year (centered to aid model convergence), a fixed effect of location, and a random effect of site nested within location. We also calculated Bayesian r-squared values for each species and combined species group (Gelman et al., 2019).

### Changes in Numerically Dominant Species

We examined shifts in the abundance of identifiable species to ask how the dominance of space by certain sessile species might have changed on rock walls over the last 42 years of environmental change. Previous work (Sebens 1985, 1986) showed that there were three historic “climax community” states on vertical surfaces in the rocky subtidal zone in the southern GOM. These climax community states were respectively dominated by *Alcyonium siderium, Aplidium glabrum,* and *Metridium senile*, all of which could retain space once occupied and not be overgrown by other species. Patches of these species persisted over many years in the same location, and sometimes mixed assemblages occurred, especially for *A. sideriu*m and *A. glabrum*. Predation by nudibranchs on anemones or soft corals could open up space, and by nudibranchs and seastars feeding on the tunicates. Invasive ascidians became common during the 42 years of study, and two of these, *Botrylloides violaceus* and *Didemnum vexillum* were of similar thickness to *A. glabrum* and were observed overgrowing it and many of the other species in the community. In addition to rate and direction of change for each species and group in the study, we also considered the potential competitive dominant species as well as the numerically dominant components of the community, and how these changes could affect the dominance hierarchy as well.

### Average Thermal Maxima

Many of the species in our study occupy a broad depth range across their geographical ranges; we selected data representing the maximum temperature encountered at each species minimum bottom depth, hereafter referred to as the average thermal maxima. We used data from the Ocean Biodiversity Information System (OBIS 2025) and temperature information from Bio-ORACLE (Assis et al., 2024) to determine each species’ average thermal maxima temperature. We used the average thermal maxima because this layer represents a more conservative estimate of the maximum temperature experienced by these organisms thereby approximating the upper limits of the fundamental thermal niche of the organism (Fuller et al., 2010). These organisms may have different thermal requirements throughout the year, but we looked specifically at their abundance during summer months in the northern hemisphere, making this the most appropriate metric of those provided by the OBIS dataset. For this analysis we excluded organisms that we could not positively identify to species and had to assign to broader groups, such as encrusting bryozoans, erect bryozoans, or hydroids.

Additionally, when species had only one known location in OBIS, we excluded them from the analysis as this would likely bias the average thermal maxima calculation. Similarly, we found confounding data reported in OBIS for *Alcyonium siderium* as only being found in the English Channel. We believe that, for New England sites, *A. siderium* has been misidentified as *A. digitatum* and therefore we used *A. digitatum data (OBIS) from this region* to identify the thermal range of *A. siderium*.

We then established temperature thresholds to identify species as cold-affinity, cool-affinity, and warm-affinity. Cold-affinity species were any with an average thermal maximum at or below our current mean summer temperature over the period from 2015-2020. Cool-affinity species had an average thermal maximum above our current mean summer temperature but below or equal to the 95th quantile of our tidbit temperature sensors recordings during the hottest months (August and September). Warm-affinity species had an average thermal maximum above the 95th quantile of our tidbit temperature sensors.

### Comparing Change in Abundance to Species Average Thermal Maxima

We wanted to evaluate the relationship between change in abundance and the average thermal maxima of species on subtidal rock walls in the GOM; we thus evaluated the relationship between their coefficient of change from the ordered beta regression models to species average thermal maxima derived from environmental occupancy as described above. We used species occurrences and environmental data layers to determine occupancy derived thermal affinities (Webb et al. 2020) to determine upper thermal maxima (termed preference) for all species sampled through the entire time series. Species occurrence records and depth distributions were taken from species occurrences in OBIS using the R (R Core Team 2024) package *robis* (Provoost and Bosch, 2017). We used these data to determine a minimum depth of occurrence for each species. We then extracted the maximum temperature, recorded over monthly climatologies from 2002-2014, at that minimum depth for each species from Bio-ORACLE (Assis et al., 2024; Tyberghein et al., 2012) using the R package *sdmpredictors* (Bosch and Fernandez, 2023; Bosch 2018).

We assessed the association between change in abundance and average thermal maxima using a linear regression with average thermal maxima predicting coefficient of change. Theory predicts that warmer preference species should increase on average while cooler preference species should decline on average with a great deal of variation in intermediate levels of average thermal maxima. We assessed model assumptions (overlap of predicted and observed distributions, linearity, homogeneity of variance, outliers) using the *performance* package (Lüdecke et al., 2021) in R and found no violations of assumptions. Finally, we evaluated the association using alternate metrics for average thermal maxima – the average maximum temperature at the mean depth these organisms are found at globally, and the greatest maximum temperature at the minimum depth these organisms are found at globally– to establish the robustness of our results. We found no qualitative differences to using average thermal maxima at minimum bottom depth (Supp. Material Table 2). Again, for this analysis we excluded organisms that we could not identify to species.

## Results

### Temperature Change in the Southern GOM

Average summer temperatures increased by more than 3 degrees from the period of 1980-1985 and 2015-2020. In the period of 1980-1985, the average summer temperature was 10.72°C, while in the period of 2015-2020 it had increased to 14.07°C (Figure 2). Over the 42 years of this study, average summer temperatures increased by 0.082°C/year with a R² = 0.46.

**Figure 2.**
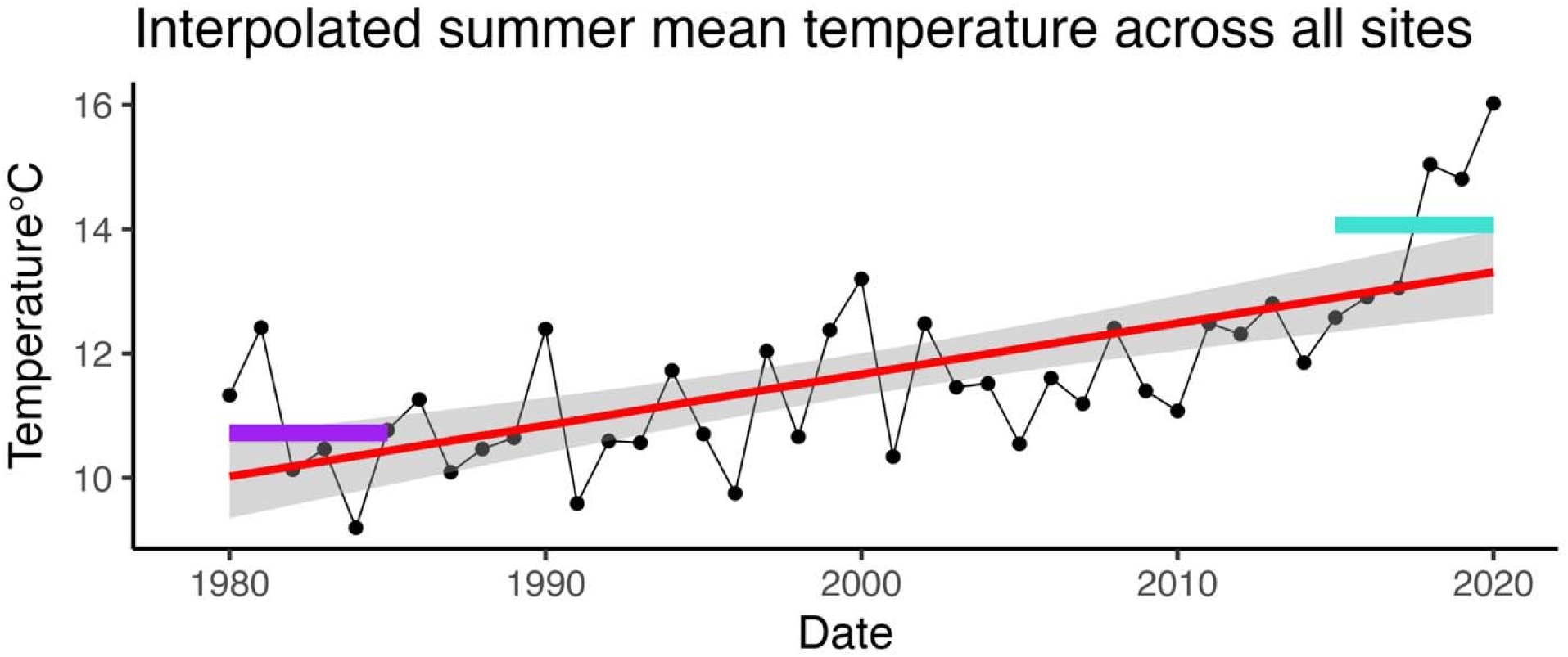
Temperature logger (Onset tidbit) data from our shallow sites (GODAS_15m interpolated), as annual summer mean temperature averaged across all 5 sites, average depth 11m. Grey shadow = 95% confidence interval. The purple line shows the average summer temperature (10.72°C) from 1980-1985, before rapid warming in the GOM was detected. The turquoise line shows the average summer temperature (14.07°C) after warming in the GOM was detected (2015-2020).

### Change in Species Abundances

Roughly equal numbers of species increased and decreased over the 42 years of the study (Figure 3). The largest coefficients of increase over time were seen among the species *Didemnum vexillum* (0.18± 0.02 SE) *Mytilus edulis* (0.17± 0.02 SE), *Edwardsiella lineata* (0.12± 0.01 SE), and *Anomia simplex* (0.12± 0.02 SE), while the greatest coefficients of decrease were seen among the species *Metridium senile* (−0.07± 0.01 SE), *Dendrodoa carnea* (−0.06± 0.01 SE), *Alcyonium siderium* (−0.06± 0.01 SE), and *Aplidium glabrum* (−0.04± 0.004 SE) (Figure 3). The confidence intervals shown in Figure 3, indicate that several species either did not change significantly or we have low precision in their estimates of change.

**Figure 3.**
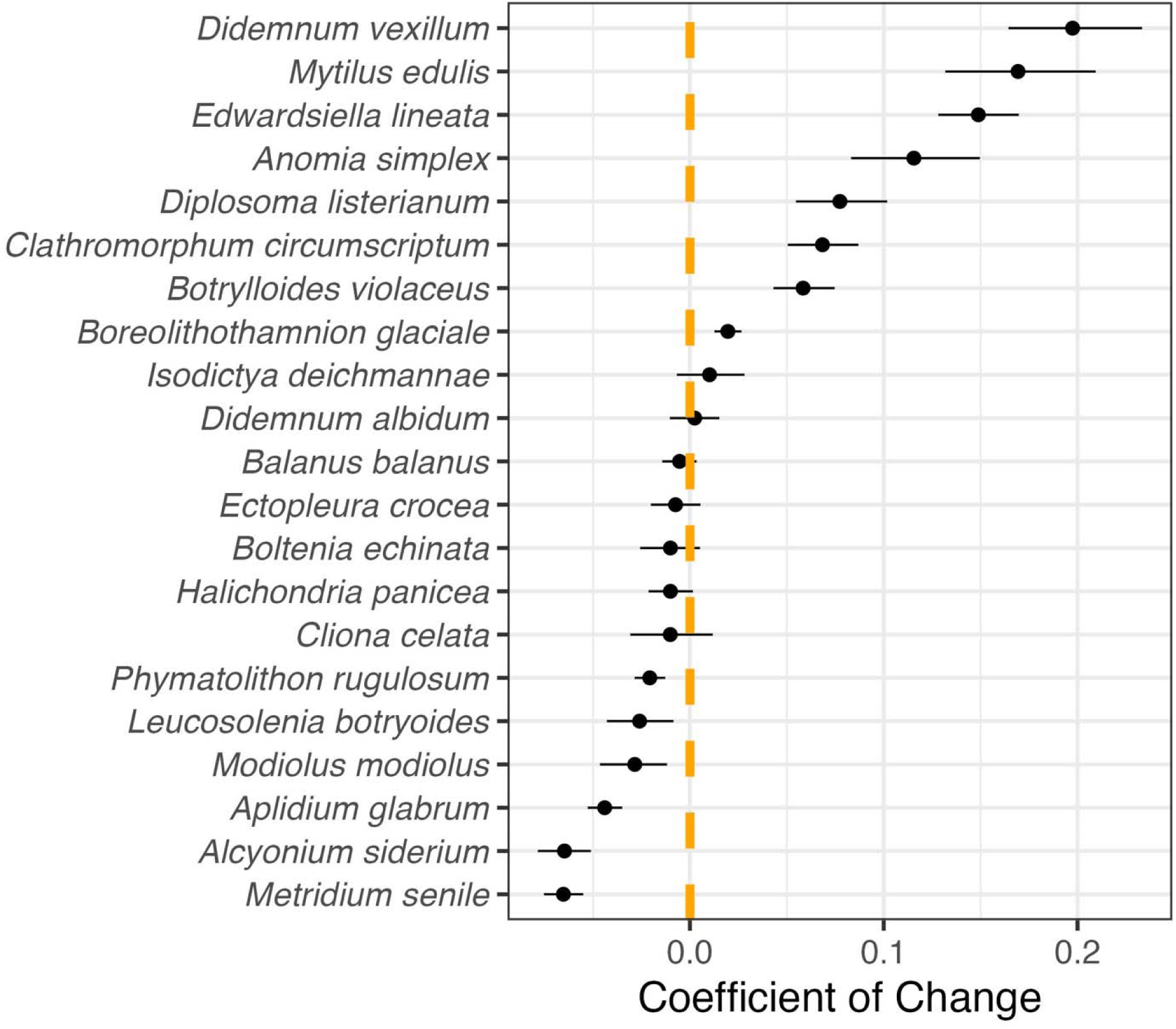
Logit coefficients of the effect of time on species abundance over 42 years of monitoring. The dashed orange line represents no change (coefficient of 0). Species to the right of the dashed line have increased in abundance while species to the left have decreased in abundance. Horizontal lines show confidence intervals.

### Changes in Numerically Dominant Species

The three numerically dominant species from the 1970s and 1980s have declined over time, some to near 0 (*Alcyonium siderium:* -0.06 ± 0.01 SE, *Aplidium glabrum:* -0.04 ± 0.004 SE, *Metridium senile:* -0.07 ± 0.01 SE). The soft coral *A. siderium,* had between 20-40% cover at these sites during the early 1980s but declined precipitously in the mid to late 1980s (Figure 4, Supp. Material Figure 1A), while staying below 10% cover at all sites where it has still been recorded through the 2020s. The original decline followed intense predation by a non-native nudibranch that first appeared here in the early 1980s (Allmon and Sebens 1988). The colonial ascidian, *A. glabrum*, declined in the mid-late 1980’s as well, falling from a peak of 75% cover at some subsites in the early 1980s to 50% cover in the mid 1980s to below 25% where it has remained at all sites throughout the period of the mid-1990s to the present (Figure 4, Supp. Material Figure 1C). A slow decline of the stalked plumose anemone, *M. senile*, followed from a peak of nearly 80% cover at some subsites, particularly those at Halfway Rock Inner and Shag Rocks Outer sites in the early-1990s down to below 25% from the mid-2000s to the present (Figure 4, Supp. Material Figure 1W). Predators on *M. senile* and *A. glabrum* were present during the entire study, and there was no observed increase in predation during these decades.

**Figure 4.**
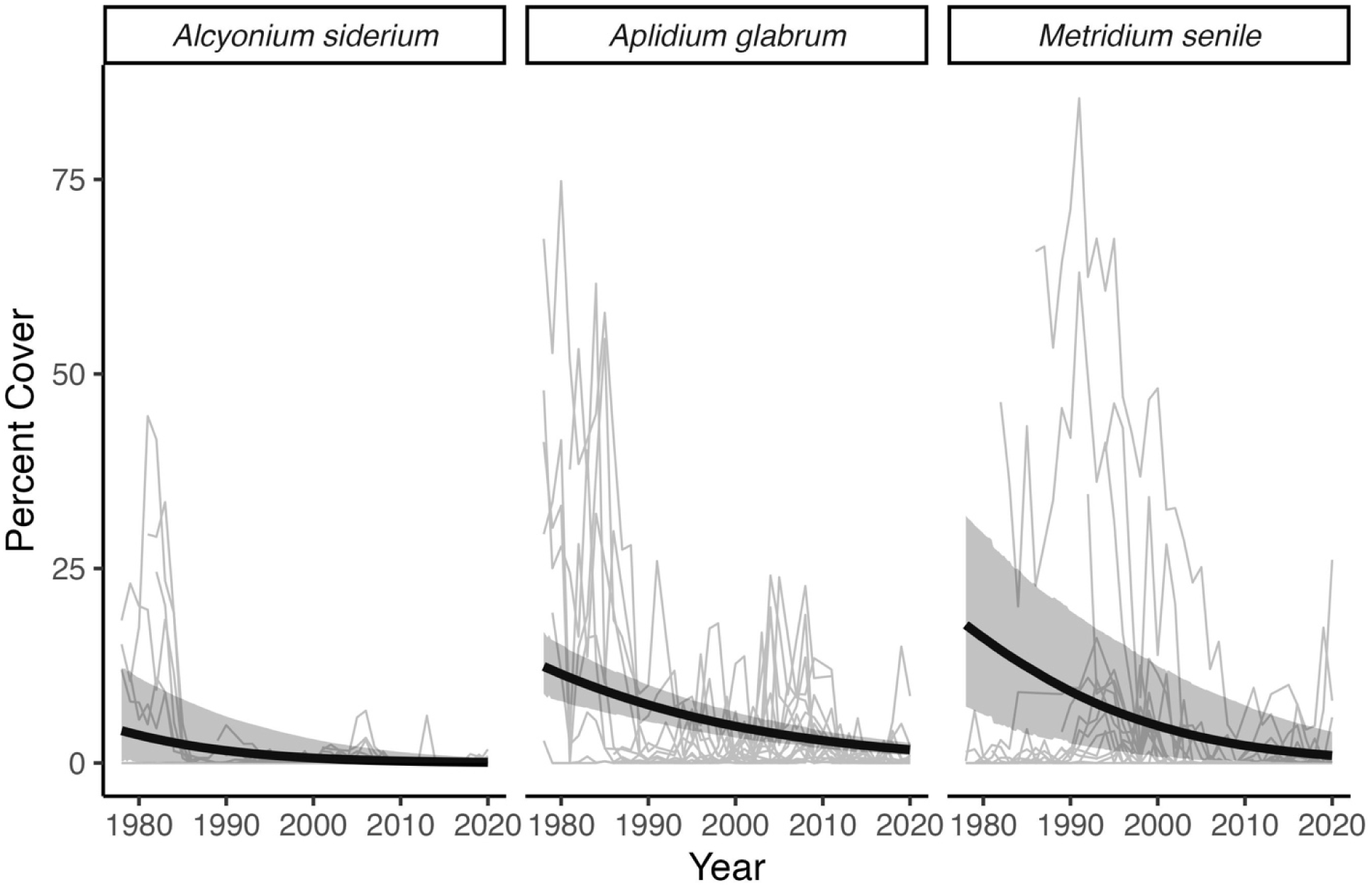
Changes in percent cover of the three numerical (and competitive) dominant species on rock walls in the GOM, *A. siderieum, A. glabrum,* and *M. senile*. Thin grey lines represent the 15 subsites surveyed. Black lines shows the trends over 42 years. Grey shadows represent 95% confidence intervals.

Several species have also increased and become highly abundant over the 42 years of this study even if rare or absent at its inception. We see the arrival of *Didemnum vexillum* (0.18 ± 0.02 SE) around 2000 (Figure 5, Supp. Material Figure 1K). *D. vexillum* was slow to gain ground at most of the sites until the early 2010s when its population exploded covering nearly 100% of some subsites around Nahant (Figure 5, Supp.

**Figure 5.**
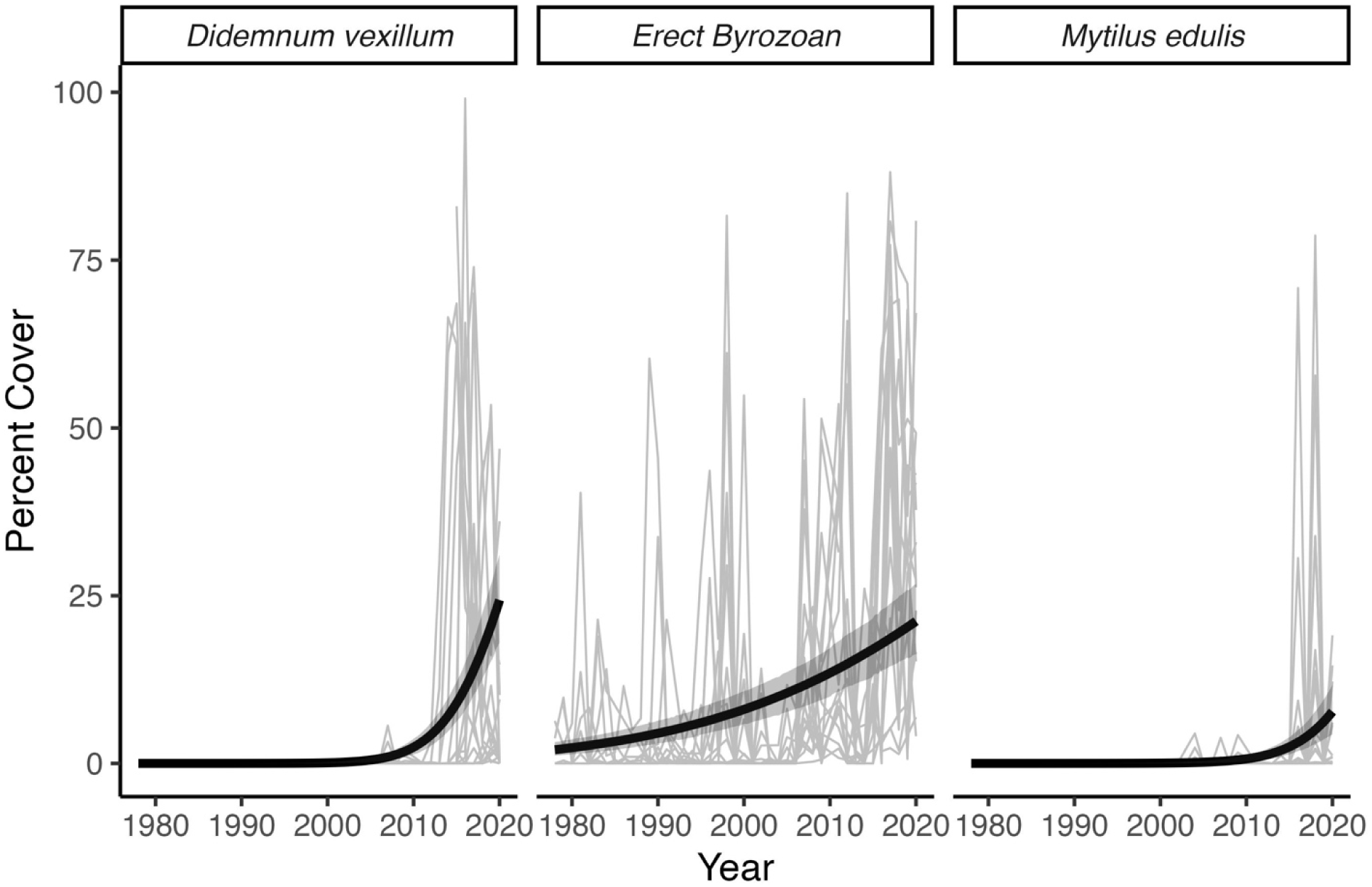
Changes in percent cover among newly dominant species on rock walls in the GOM. Thin grey lines represent the 15 subsites surveyed. Black line shows the trend over 42 years. Grey shadow represents confidence interval.

Material Figure 1K). The multi-species conglomeration of erect bryozoans (0.06 ± 0.005 SE) covered less than 20% of walls through the late 1980s before increasing to peak coverage of over 75% in some places during the 2000s, while maintaining around 50% coverage at most subsites (Figure 5, Supp. Material Figure 1P). The erect bryozoan complex was maintained at a low abundance through the late 1980s (Figure 5), but showed cyclical peaks in abundance every decade following, with abundance near 75% coverage at some sites through 2020 (Figure 5, Supp. Material Figure 1P). A pulse recruitment event of the blue mussel *M. edulis* (0.17 ± 0.02 SE) occurred in the mid 2010s with coverage nearing 75% at one subsite in Nahant (Figure 5, Supp. Material Figure 1Y). A second smaller *M. edulis* recruitment event occurred in the late 2010s at multiple subsites in Nahant (Figure 5, Supp. Material Figure 1Y).

### Average Thermal Maxima

We found that this group of species varies greatly in their average thermal maxima. Broadly, we see that species in this community prefer (i.e. occupy regions with) temperatures below the 95th quantile of summer temperatures, which was 17.4°C, and 4 species prefer temperatures below 14.07°C, our mean summer temperature between 2015-2020 (Figures 6 and 7). Species that have an average thermal maximum above 17.4°C include the non-native colonial tunicates *Diplosoma listerianum* and *Botryloides violaceus,* the tubular hydroid *Ectopleura crocea,* the sponge *Isodictya deichmannae,* and the mussel *Mytilus edulis. Edwardsiella lineata*, the lined anemone and *Anomia simplex*, the common jingle shell, each have an average thermal maxima near 25°C (Figures 6 and 7).

**Figure 6.**
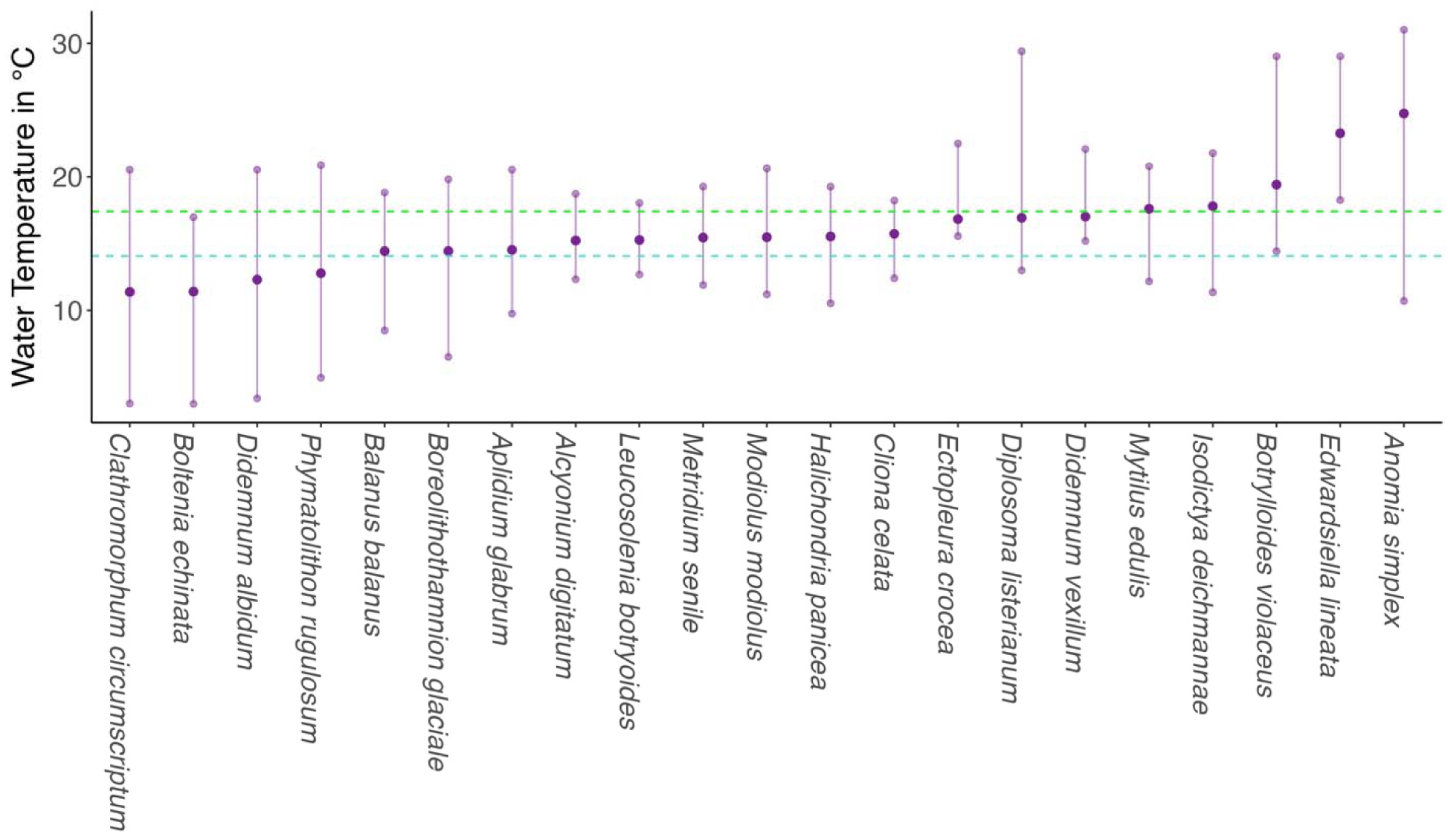
The maximum temperature experienced by species at our locations at their minimum global depth. Dark purple circles represent the mean maximum temperature found at their global minimum depth while purple bars represent the range (maximum-minimum) of temperatures experienced at their minimum global depth (data from OBIS). The green dashed line depicts 17.4°C while the turquoise dashed line represents 14.07°C.

**Figure 7.**
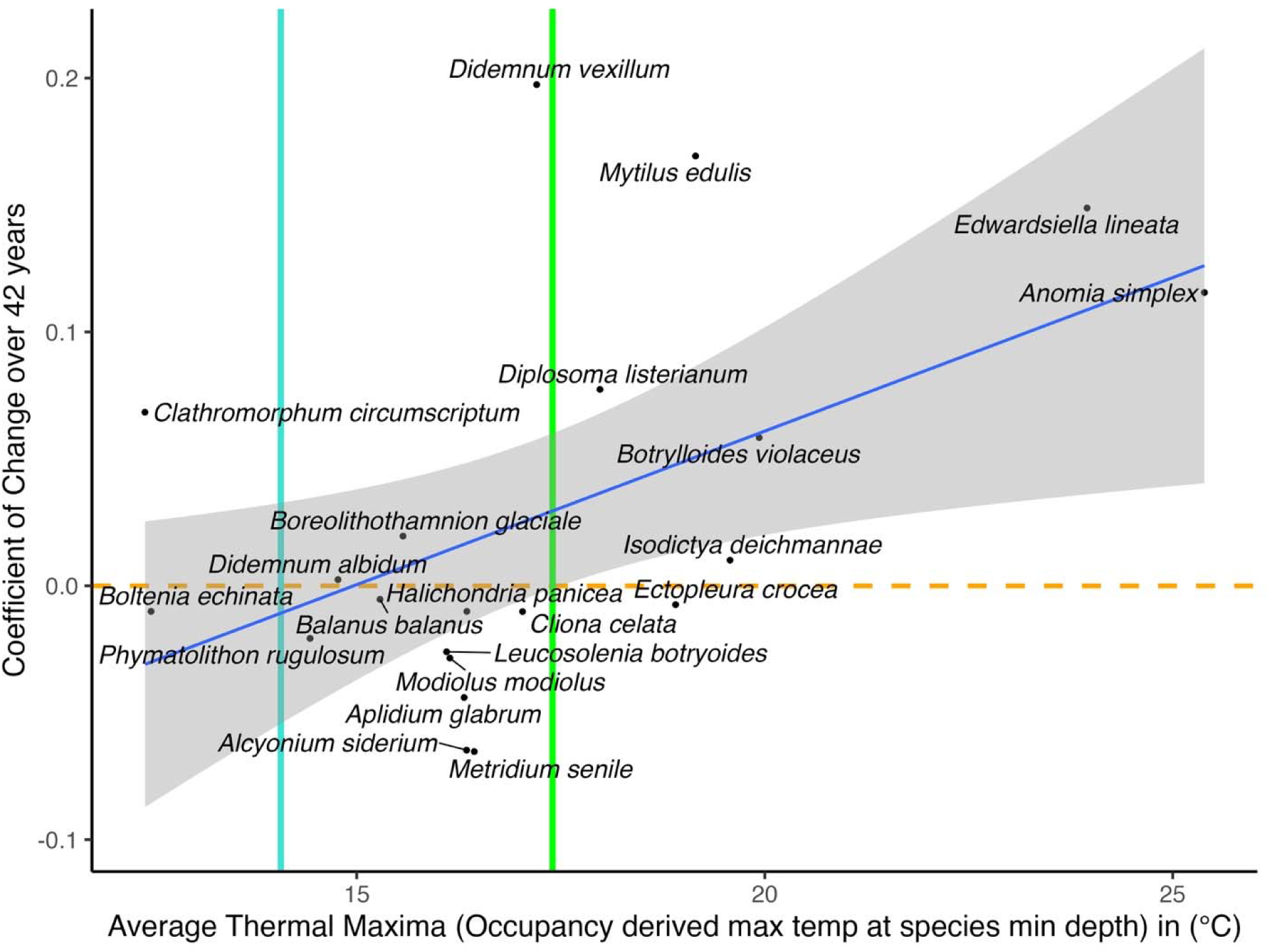
The average thermal maxima of species found at the two locations surveyed paired with their change in abundance over 42 years. The blue line (R² = 0.26) shows a positive slope indicating that warm-affinity species have been increasing. The orange dashed line marks zero change in abundance over the period of this study. The turquoise line separates the cold-affinity species (<14.07°C) from the cool-affinity species (14.07-17.4°C). The green line at 17.4°C separates the cool-affinity species (14.07-17.4°C) from the warm-affinity species (>17.4°C)

### Comparing Change in Abundance to Species Average Thermal Maxima

There is a positive relationship between average thermal maxima and change in abundance (P value = 0.018, R² = 0.26). The species that increased the most are associated with warmer temperatures (Figure 7). Species that decreased in abundance all inhabit locations with maximum summer temperatures at or below 17.4°C, including *Alcyonium siderium, Metridium senile* and *Aplidium glabrum*, the horse mussel *Modiolus modiolus,* and the asconoid sponge *Leucosolenia botryloides* (Figure 7). Seven species did not change in abundance over the period they were monitored (Figure 3). This includes 1 cold-affinity species (*Boltenia echinata*), 4 cool-affinity species *(Didemnum albidum, Balanus balanus*, *Halichondria panicea*, and *Cliona celata*), and 2 warm-affinity species (*Ectopleura crocea and Isodictya deichmannae*) (Figure 7).

## Discussion

The loss and gain of species on the subtidal rock walls monitored in this study are consistent with warming trends documented for the Gulf of Maine. This represents the beginning of a substantial change in species composition through time - temporal turnover (*sensu* Pinsky et al. 2025) resulting in a major shift in community composition. Previously dominant species are now rare in this system, while newly arrived non-native species have established growing populations. There have been some “winners” (*Edwardsiella lineata*, *Anomia simplex*, *Mytilus edulis*, *Diplosoma listerianum*, *Botrylloides violaceus*, and *Didemnum vexillum*) and some “losers” (*Metridium senile, Aplidium glabrum,* and *Alcyonium siderium*), while some 7 other species show close to no change in their abundance over 42 years of monitoring (Figure 3). Species showing no change also include one cold-affinity species, *Boltenia echinata.* Such species which did not change in abundance show a remarkable ability to withstand physical and biological stressors including warming temperatures and competition with non-native species.

The warm-affinity species, which generally increased in abundance, likely predict the future makeup of rock wall communities in the southern Gulf of Maine. It has been predicted that communities will respond to changing environmental conditions by shifting towards more thermally-tolerant species while thermally sensitive species are expected to decline (Holbrook et al., 1997; Pinsky et al., 2025). We see this as a general trend among the species in our study (Figure 7) where the organisms that have increased the most in abundance are warm-affinity species. Species that appear to be thriving in the locations surveyed include the non-native tunicate *Didemnum vexillum*, the commercially and ecologically important blue mussel *Mytilus edulis*, the lined anemone *Edwarsiella lineata*, and the common jingle shell *Anomia simplex*. The latter two species can withstand a maximum temperature of or near 25°C (Figure 7). This indicates they should continue to increase in abundance in the coming years. The non-native ascidians *Botrylloides violaceus* and *Diplosoma listerianum* increased in abundance during the first half of the monitoring period, then decreased, but are still common in these communities.

Our cool-affinity species all have an average thermal maxima above the average summer temperatures at our sites of 14.07°C, however their average thermal maxima fall below the 95th quantile of summer temperatures, which was 17.4°C.

With the exception of the coralline crust *Boreolithothamnion glaciale*, all cool-affinity species declined or maintained similar abundances over the course of the study indicating that summer temperatures may prove too warm for many of them to persist at our sites (Figure 7). This has important implications for future species richness on subtidal rock walls in the GOM as many of these cool-affinity species may be lost as temperatures continue to warm. In the case of B. glaciale, it may have benefitted from loss of competitors that would normally overgrow it, and by increased sea urchin grazing on rock walls with less soft fleshy invertebrates (Sebens 1985, 1990). Our cold-affinity species include just the coralline algal crust *Clathromorphum circumscriptum* and the tunicate *Boltenia echinata*. *B. echinata* is relatively rare at our sites, and remained at a similar albeit low abundance throughout the length of the study. C. circumscriptum may also have benefitted from loss of some of the invertebrates that would normally overgrow it.

Some species can respond to warming ocean temperatures associated with climate change by shifting their populations towards the poles (Barry et al., 1995; Holbrook et al., 1997; Lubchenco et al., 1993; Pinsky et al., 2013). While this appears to be a common response of marine organisms (Barry et al., 1995; Beaugrand et al., 2002; Holbrook et al., 1997; Pinsky et al., 2013; Southward et al., 1995; Sunday et al., 2012), currents and geographical boundaries constrain the poleward dispersal of organisms with planktonic larvae (Dunstan and Bax, 2007; Gaylord and Gaines, 2000). As currents flow largely in the opposite direction of climate velocities in the coastal GOM, they are likely an impediment for most of the species in our study. Pinsky et al. (2013) shows that when poleward dispersal is limited by a geographical boundary, species tend to shift into deeper colder waters. Whether or not this shift is the case for sessile organisms on subtidal rock walls in the GOM remains to be seen, and our study sites do not include much greater depth zones.

## Conclusion

A pressing challenge in this era of rapidly changing climate is to understand species and community-wide responses to environmental change. Our ability to predict even near-future responses requires an understanding of the processes that drive community structure, and how those are affected by changing climate conditions. The rate of temperature increase in the oceans has been dramatic in areas such as the GOM. Here we have seen sea surface temperatures rise 0.23°C per year, more than twenty times the global average increase of 0.01°C per year (Pershing et al 2015), while at our sites we have seen mean summer temperatures rise from 10.72°C during the period of 1980-1985 up to 14.07°C during the period from 2015-2020 (Figure 2). Long-term studies such as this one can help elucidate whether we are in the midst of a temporal turnover of species, versus a short-term deviation that could reverse. Overall, this study found that temporal turnover is occurring on subtidal rock walls in the southern GOM, and includes some of the dominant space occupiers, and competitively dominant species that were once able to acquire and hold space for many years.

Over the last several decades the GOM has seen a wide range of biotic and abiotic changes (Byrnes et al., 2024; Lotze et al., 2022; Record et al., 2023; Steneck et al., 2013), yet still retains a diverse community of sessile invertebrates throughout its rocky subtidal zone. Historically, on vertical surfaces at our sites, this invertebrate community was initially dominated by three species, *Alcyonium siderium, Aplidium glabrum,* and *Metridium* senile that form long-lived aggregations and were able to exclude other species and persist for many years (i.e. alternate climax states, Sebens 1985), however the abundance of these species has been reduced dramatically, including nearly 10-fold for *A. siderium* (Figure 4). These populations were decimated by a non-native predator, the nudibranch *Tritonia plebeia* in the 1980s, which then disappeared from the region (Allmon and Sedens 1988). Three decades later, without that predator present, populations have never recovered and are now represented by scattered individuals.

These climax community members have seemingly been replaced by the non-native tunicate *Didemnum vexillum,* the commercially and ecologically important blue mussel *Mytilus edulis*, and an assemblage of erect bryozoans, as well as the non-native ascidian *Botrylloides violaceus* and the native anemone *Edwardsiella lineata* at certain sites and time periods. Additional declines were seen among many species that normally experience maximum temperatures below 17.4°C (Figure 7). Our results show that communities here have responded to warming temperatures by shifting their component species to ones that are more thermally-tolerant. Especially in disturbed areas, this new community assemblage may be based on non-native species, while other areas might see colonization by native species that follow climate velocities towards the poles. This ongoing long-term study offers a glimpse of what subtidal rock wall communities may become in the future as warming increases and more cold adapted species are lost from the system.

## Acknowledgements

We thank the following people who led or assisted on many field SCUBA surveys at our long-term sites over several years each, and/or helped with analysis of the resulting data: First decade: Mark Patterson, Stephen Norton, Ray Merva, Robin Aiello, Larry Lipke, Richard R.(Randy) Olson, Mark Ashenfelter, Joann Resing, Donald R. Levitan, John Sigda, David Low, Julia Miles, Doug Updike, Edward (Ted) Maney, Steve Zamojski. Second decade: C. Shannon Briscoe, Kathy Paull, Kathy Durante, Chuck Arnold, Charley Ellis, Sal Genovese, Krista Graham, Brian Helmuth, Karen Vandersall, Sean Grace. Third decade: Assaf Gordon, Brad Agius, Gretchen Goodbody, Richard Sleboda, Tom Eldridge, Nathan Geraldi, Robert Miller, Kenan Matterson, Tim Dwyer, Robin Elahi. Fourth decade on: Kevin Turner, Ted Lyman, Curtis Fahey, Christopher D. Wells, and Angela Jones. We also thank the numerous students and other assistants who worked on the quantitative analysis of quadrat photographs (1980-2020), especially Melody Chen, Kelton Clark, Ali Rhoades, Autumn Turner, Nicki Le Baron, Alifaire Noreen, Audrey Olshefsky, Fiona Gettinger, Irina Gettinger and Robin Gropp, and the many other divers who assisted for short periods during all decades.

## Notes

### Competing Interest Statement

The authors have declared no competing interest.

