## Supplementary Data for "Changes in species composition of sessile communities on subtidal rock walls in the southern Gulf of Maine during four decades of warming"

Table 1 Sites and subsites photographed for analyses, with date the subsite was established and approximate depth of the subsite. Each subsite is approximately	0.3-0.5 m² in the center of a vertical surface approximately 1-2 m^2^.

**Table 1. Sites and Subsites**

Site	Subsite Names	Initial Sample	Approximate depth (m)

HRI	WA (Wall Away)	April 1979	16

VC (Vertical Control)	April 1979	16

VS (Vertical Shallow)	April 1979	7

HRO	VCT (Vertical Control Top)	September 1981	9

VCB (Vertical Control Bottom)	September 1981	10

VDL (Vertical Deep Left)	May 1989	15

VDR (Vertical Deep Right)	May 1989	15

SHI	WA (Wall Away)	September 1978	8

VC (Vertical Control)	August 1978	8

SHO	VC (Vertical Control)	December 1985	9

VS (Vertical Shallow)	February 1986	8

V3 (Vertical 3)	September 2000	8

DB	WA (Wall Away)	October 1978	7

VC (Vertical Control)	April 1978	7

ALC (Alcyonium Wall)	February 1988	7

Table 2 Comparison of BIOracle temperature layers as alternate metrics for average thermal maxima.

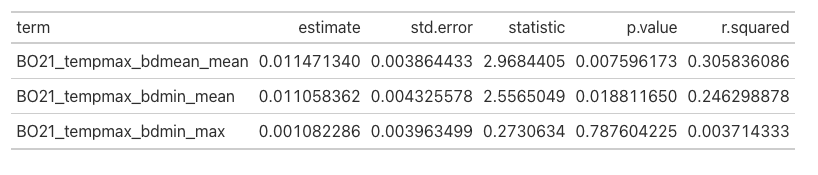

Supplemental Figure 1. Percent Cover of all organisms identified in this study at each of the 5 sites.

A.

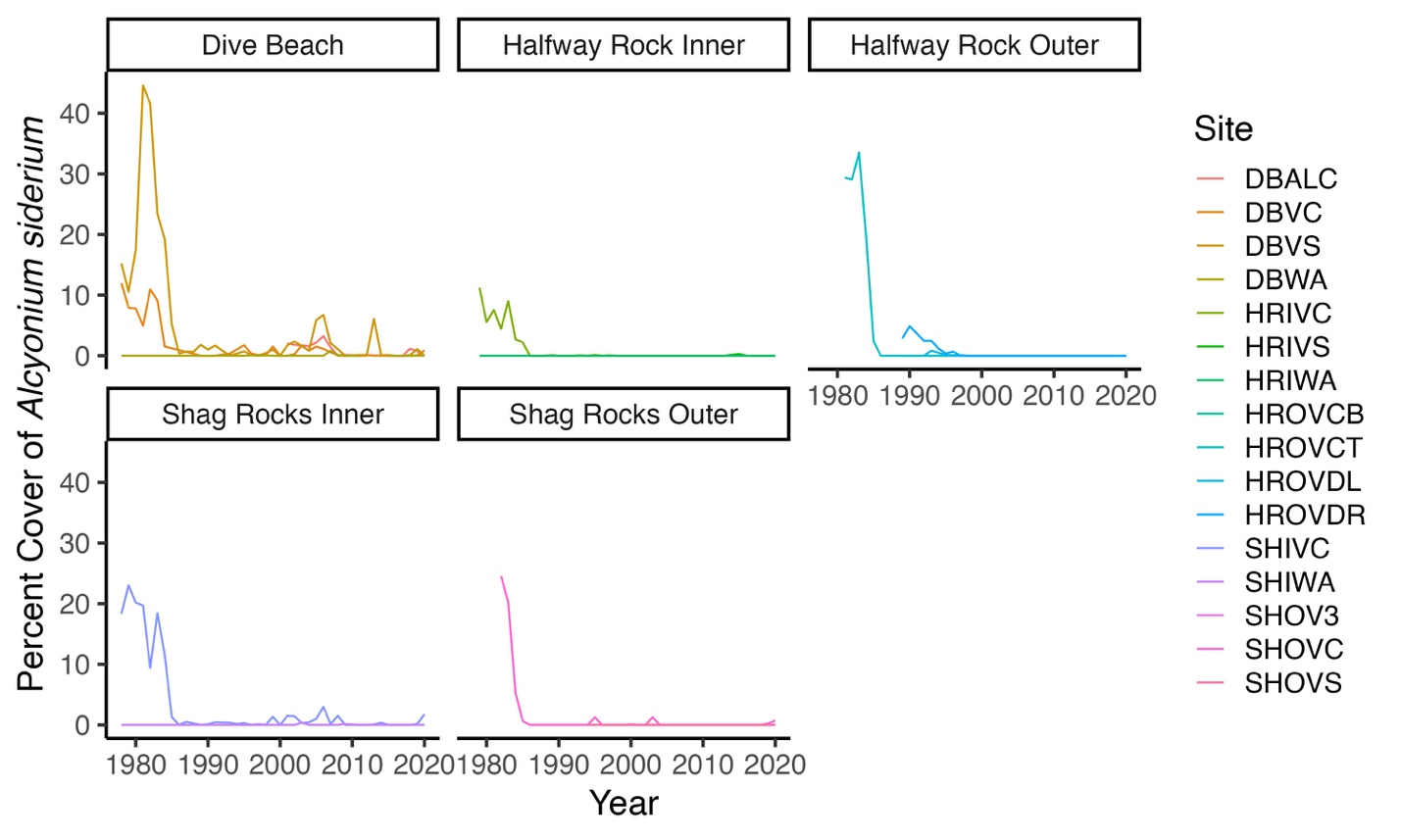

B.

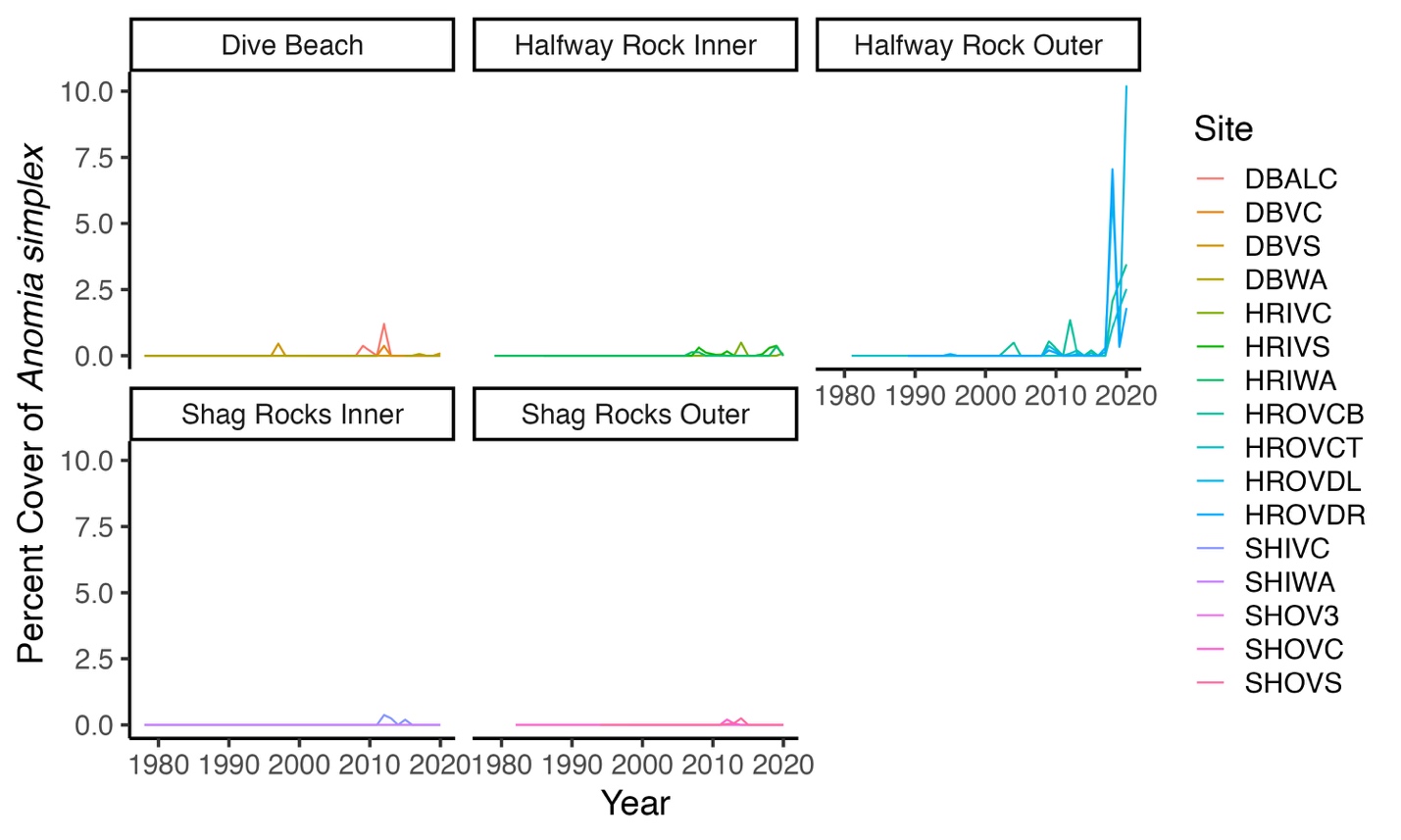

C.

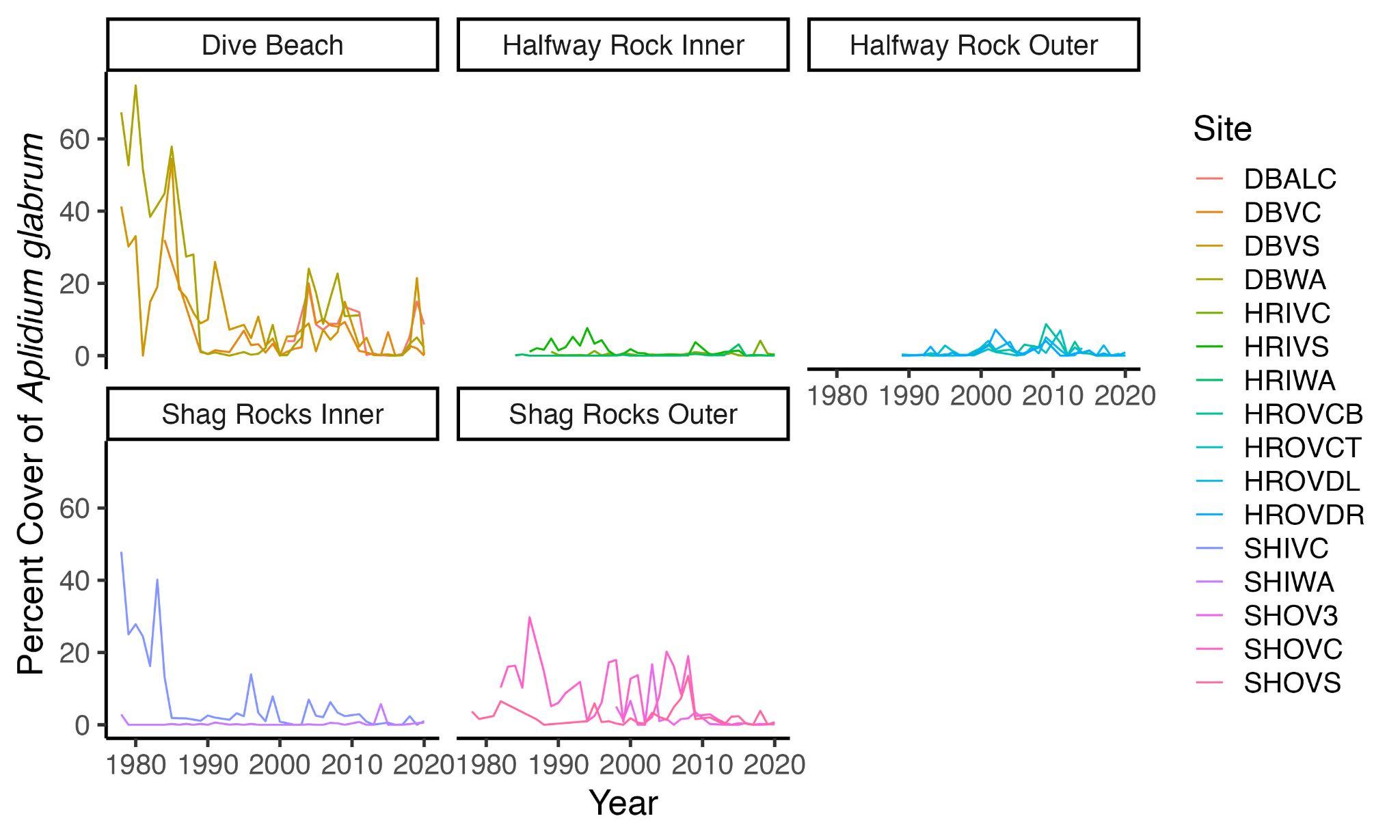

D.

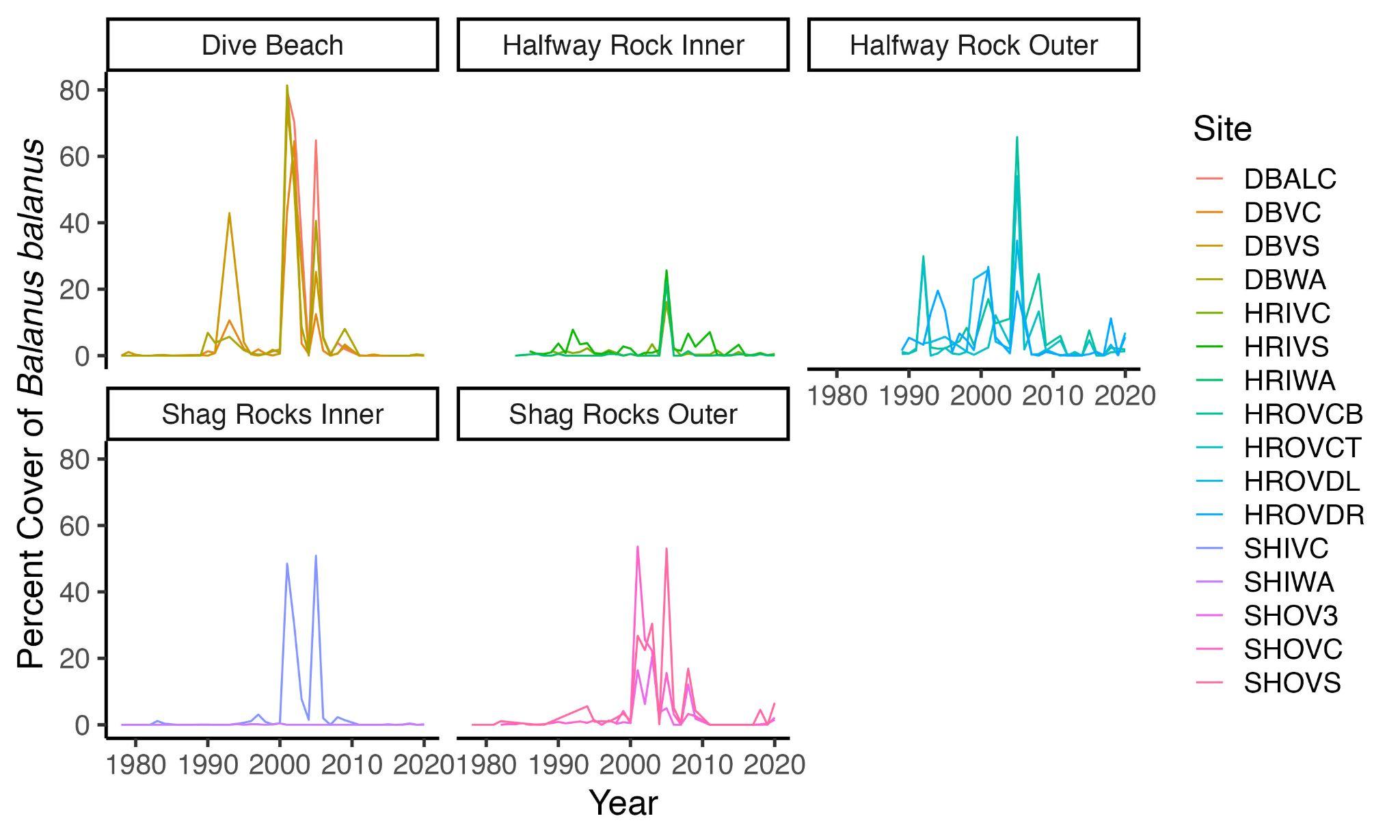

E.

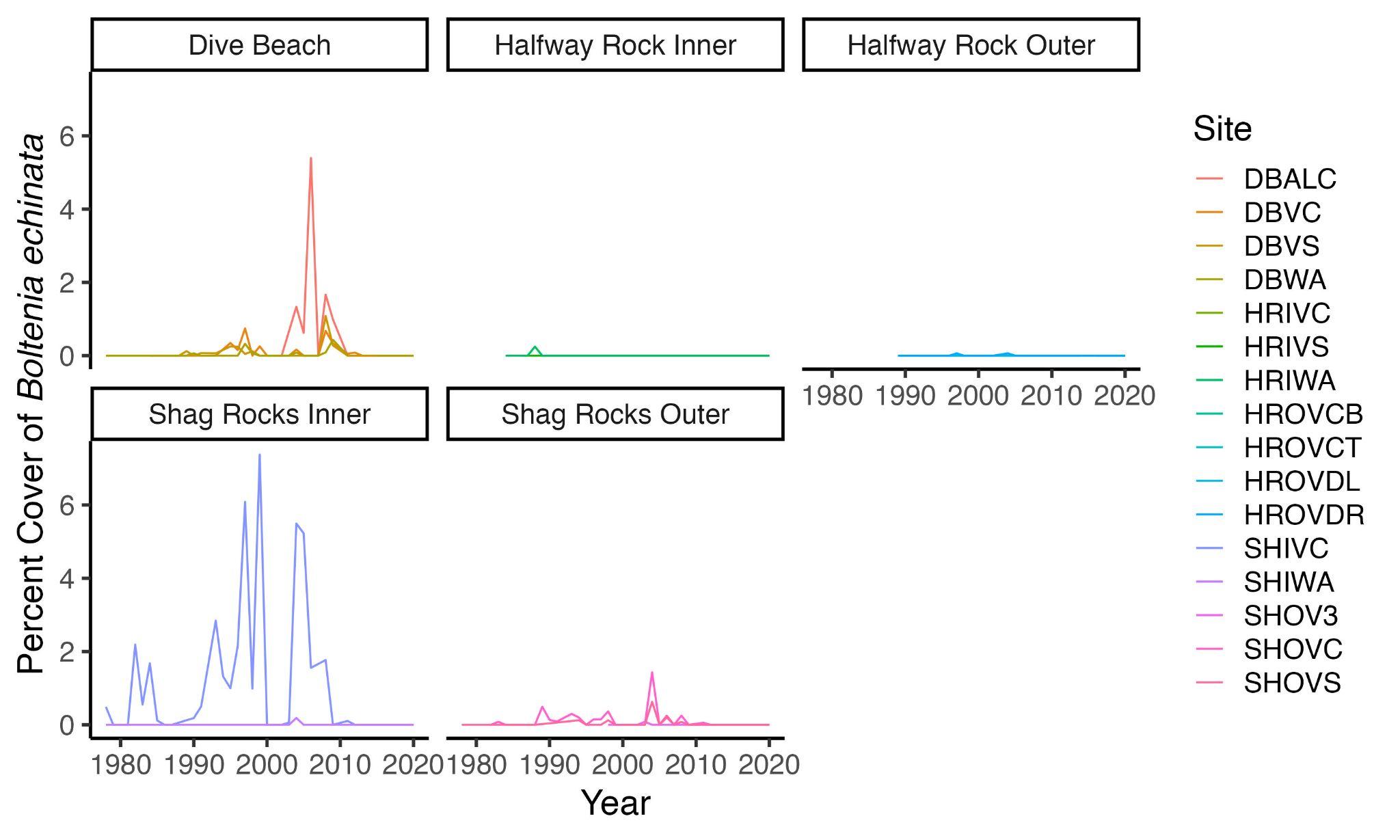

F.

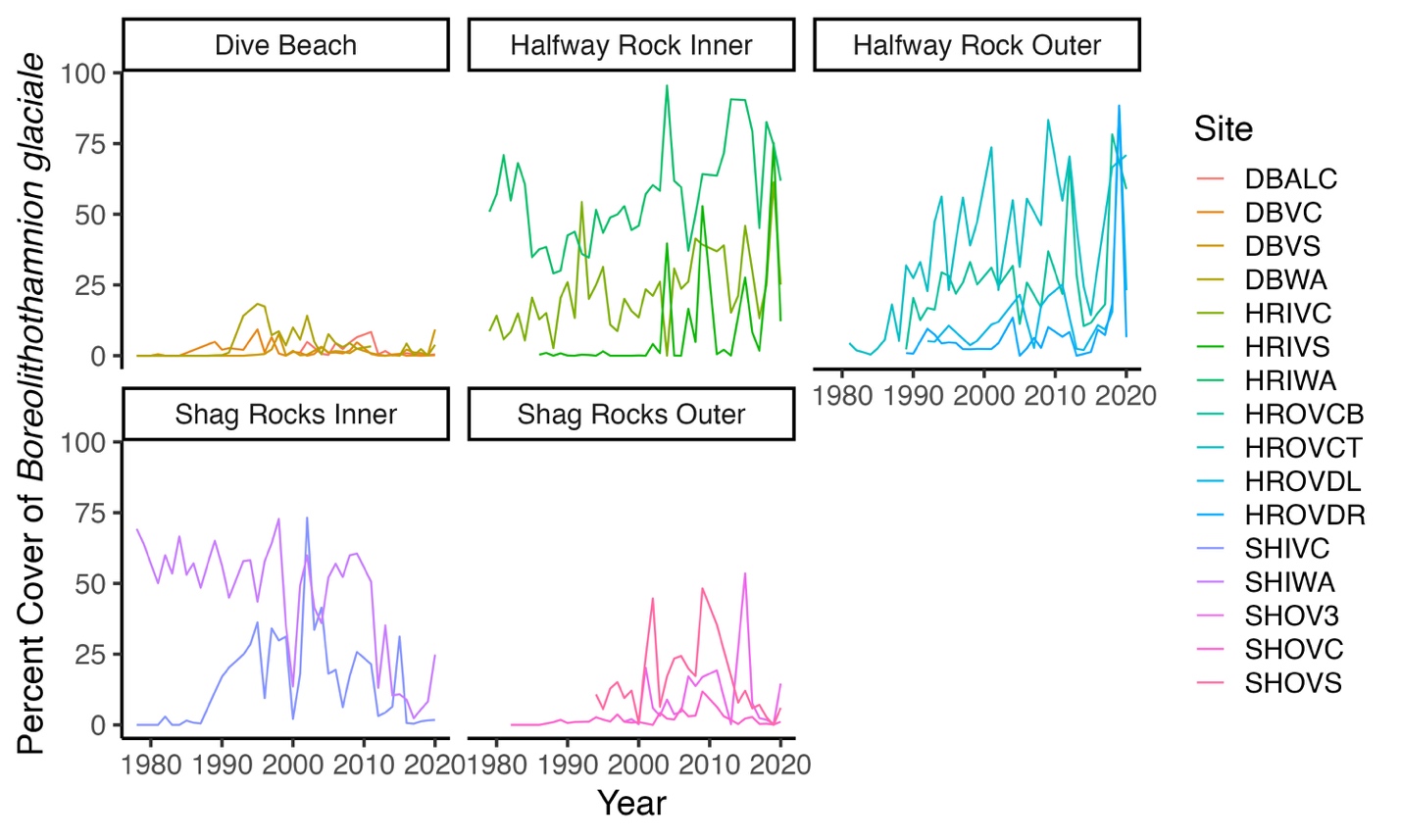

G.

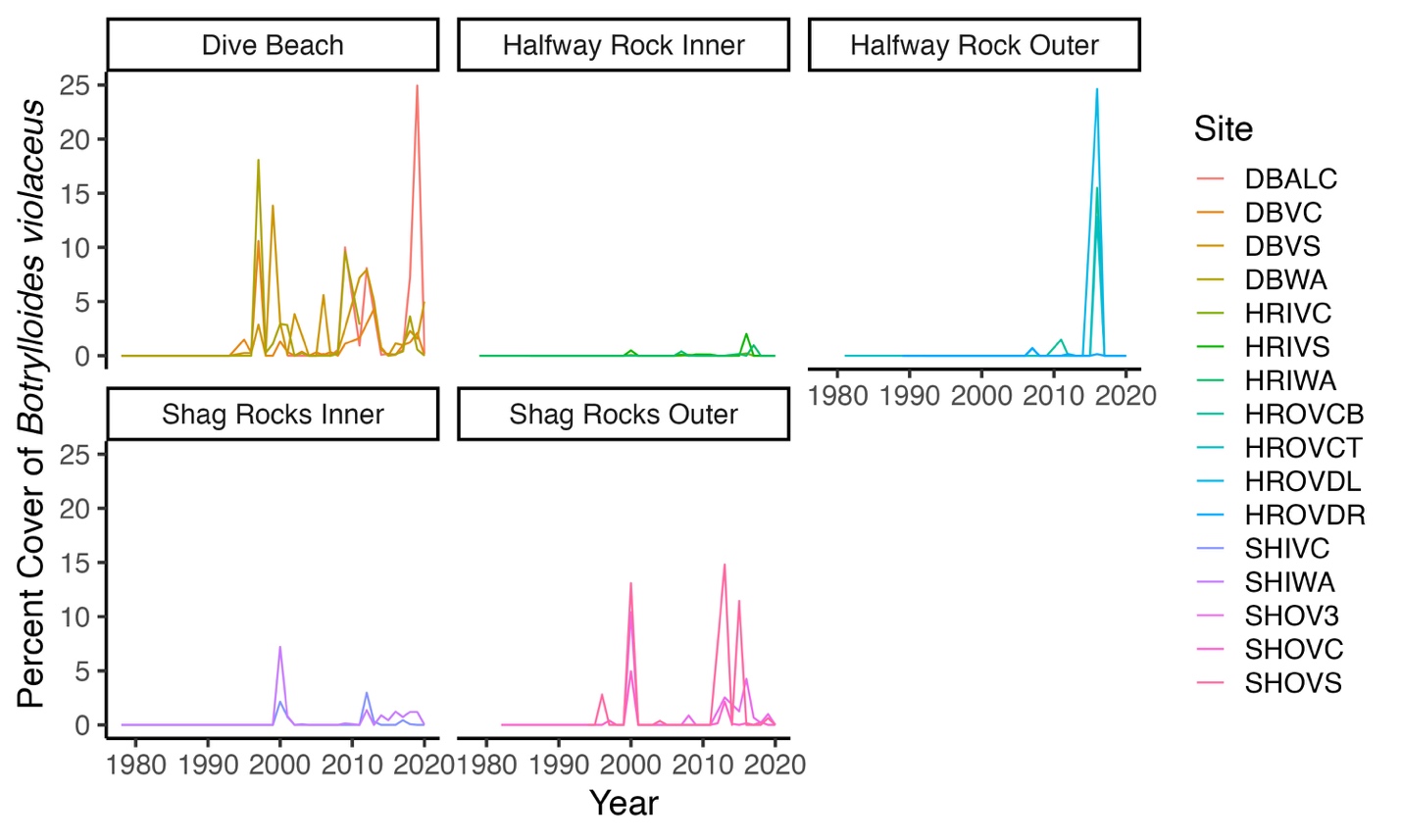

H.

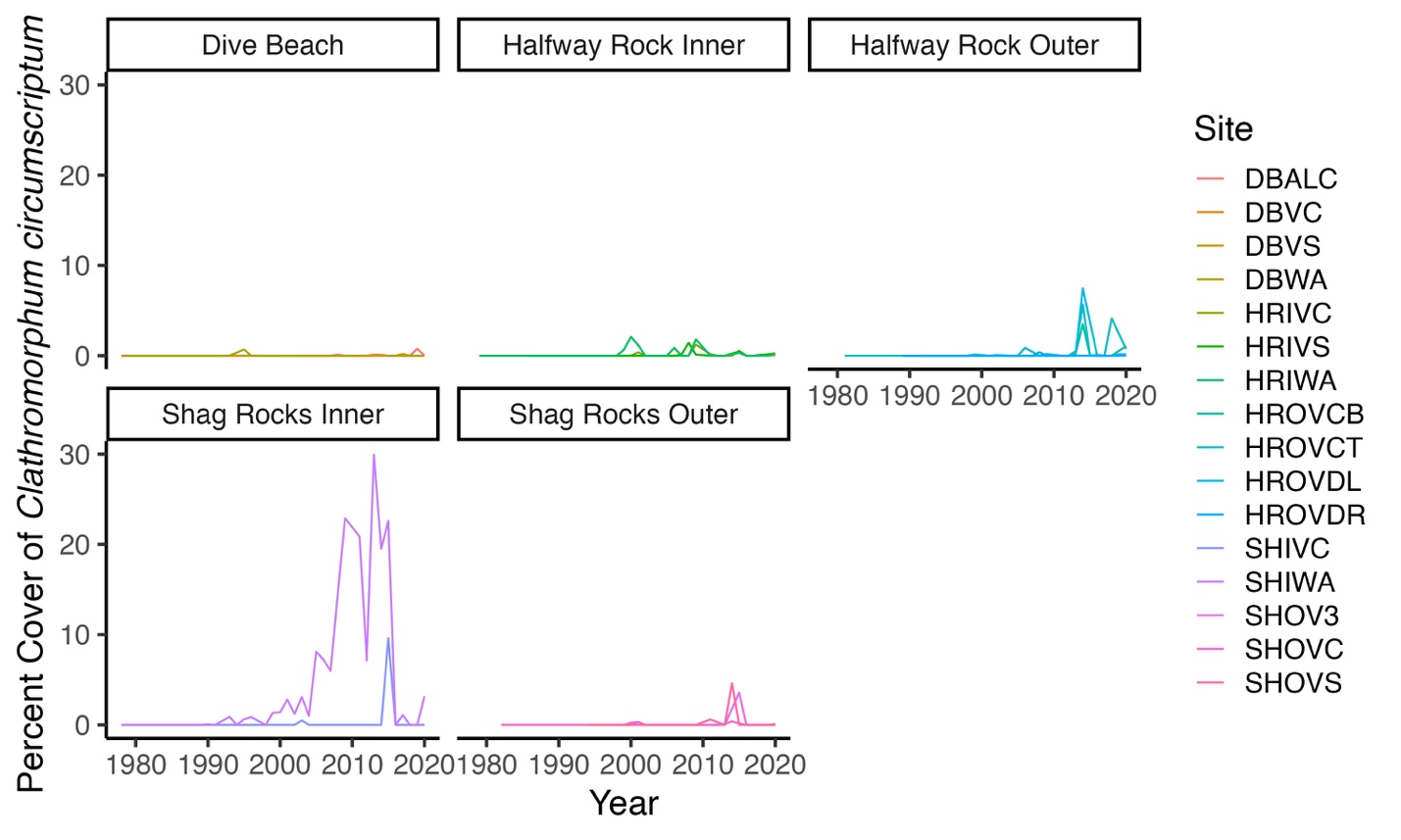

I.

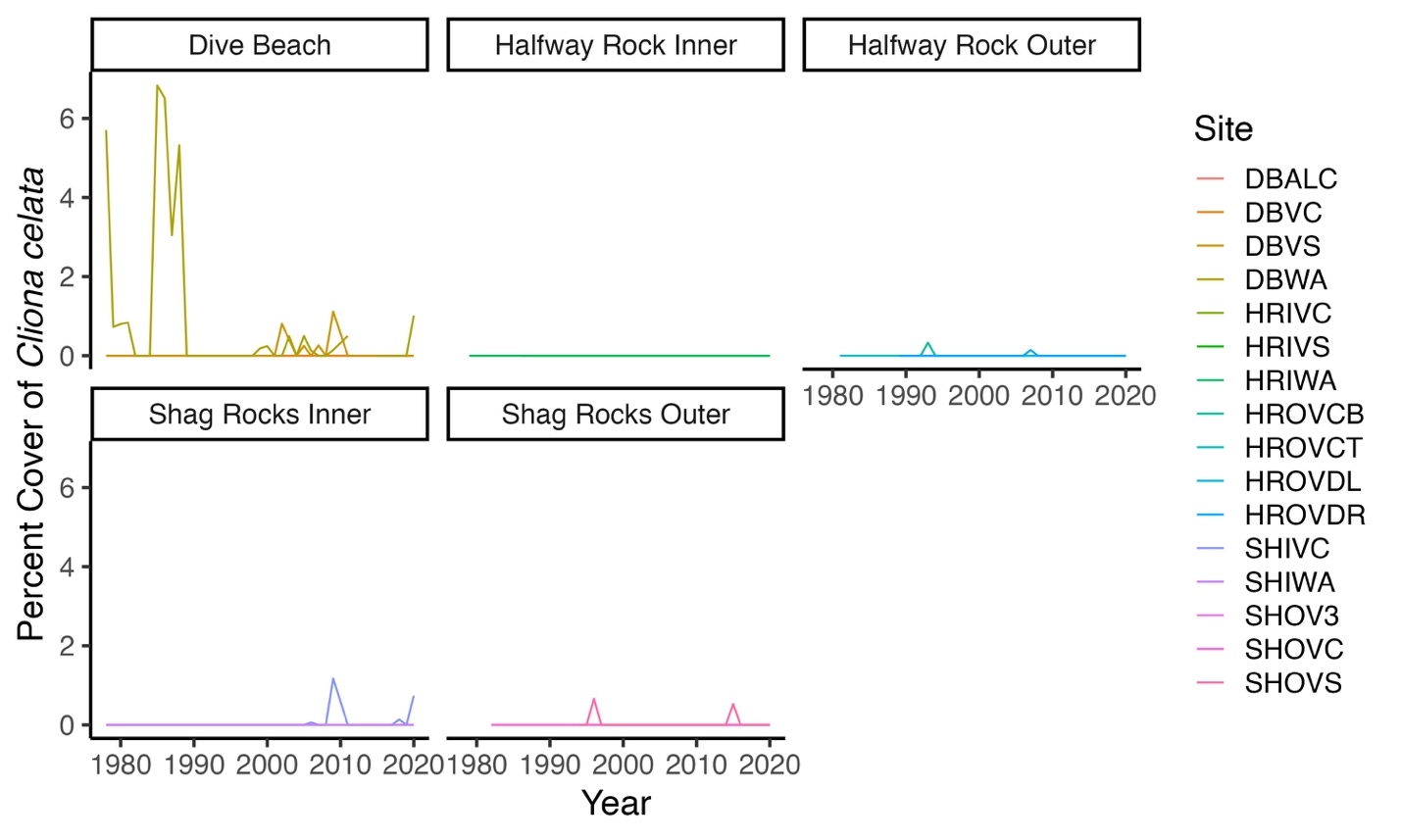

J.

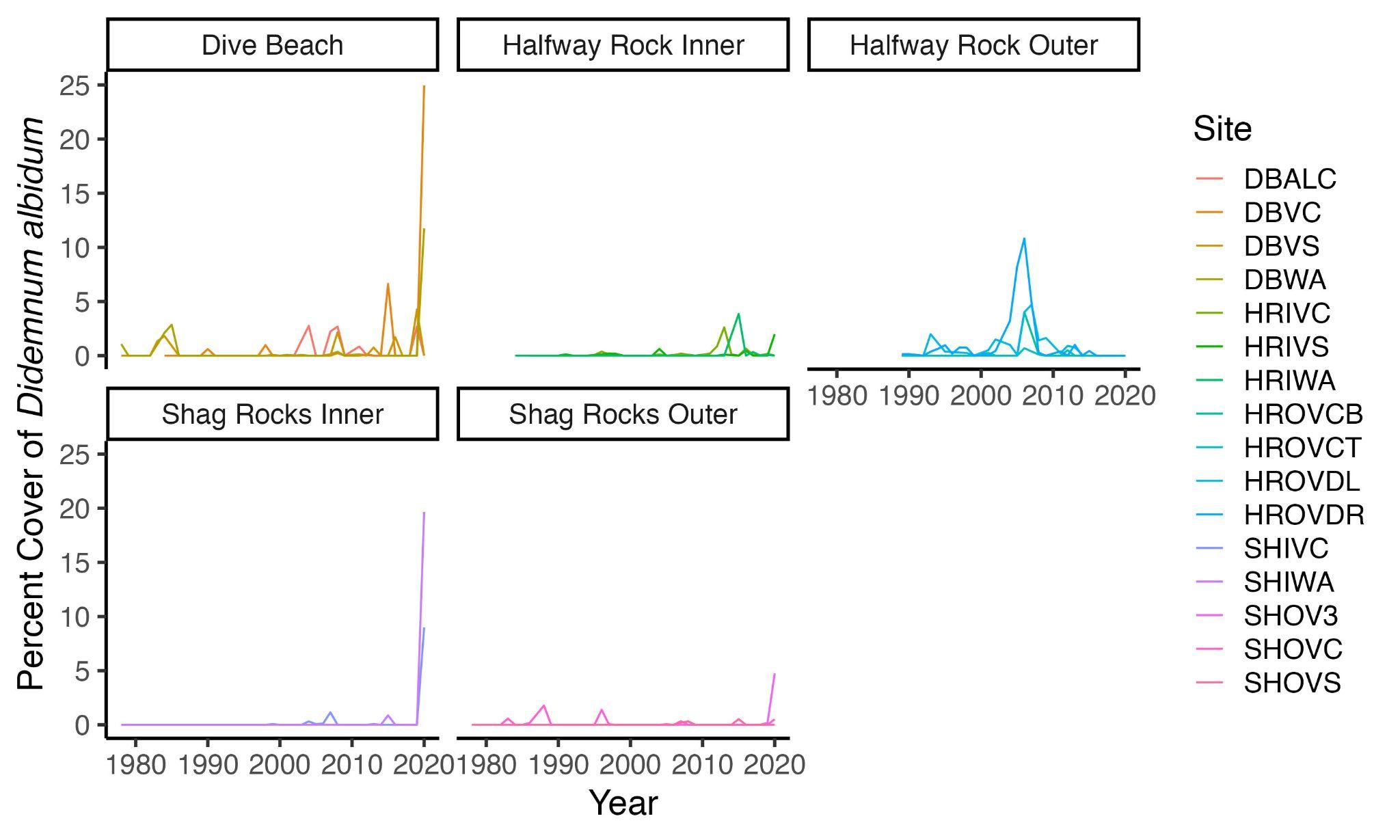

K.

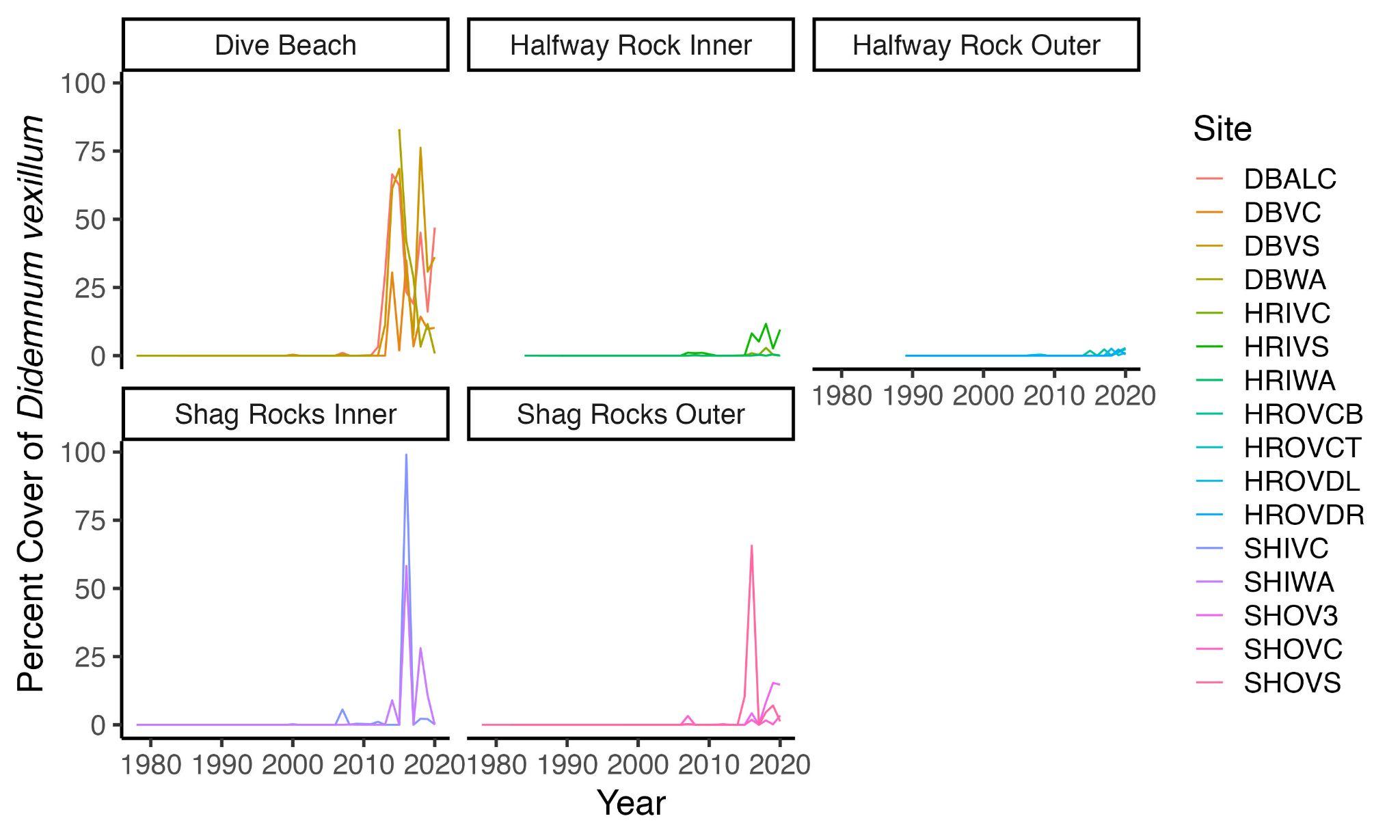

L.

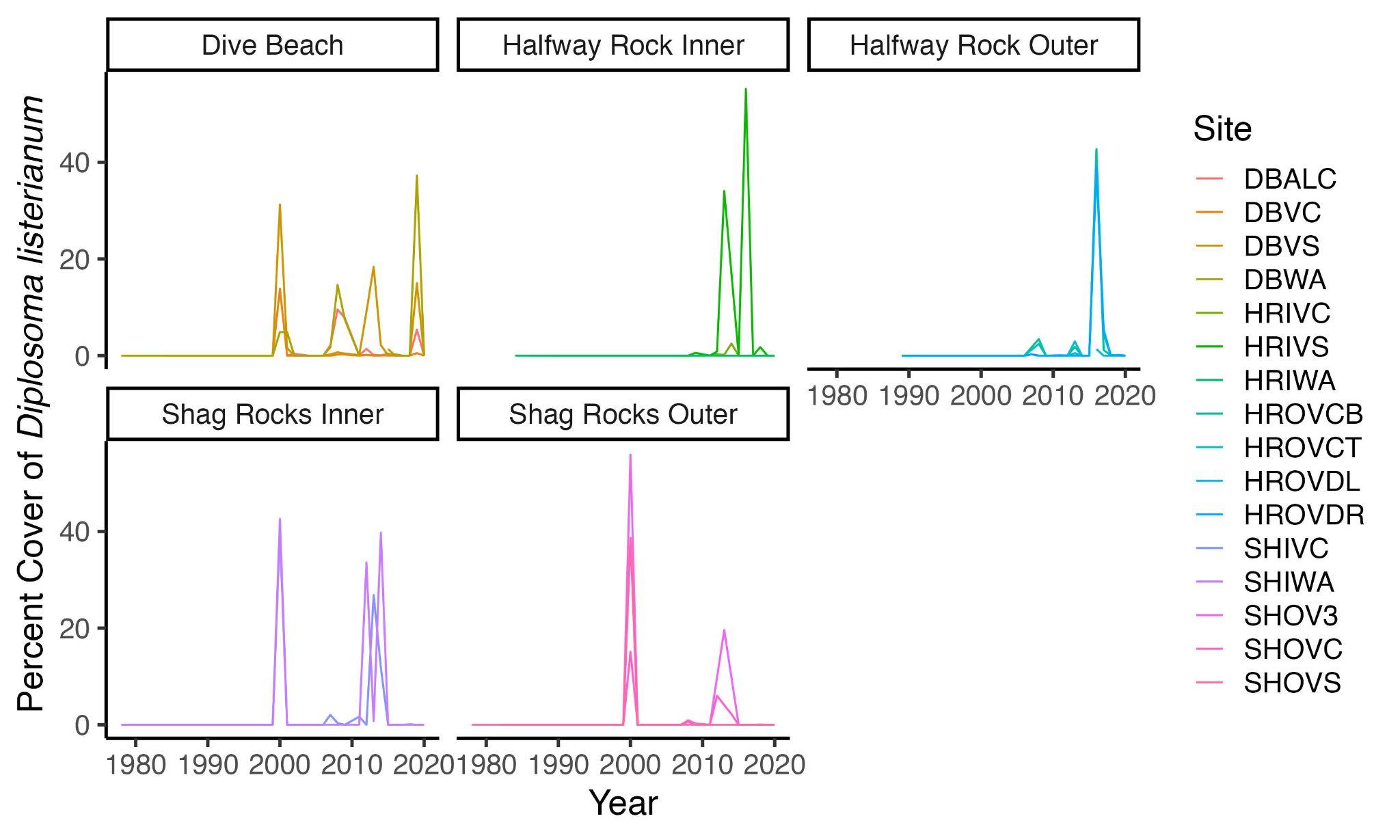

M.

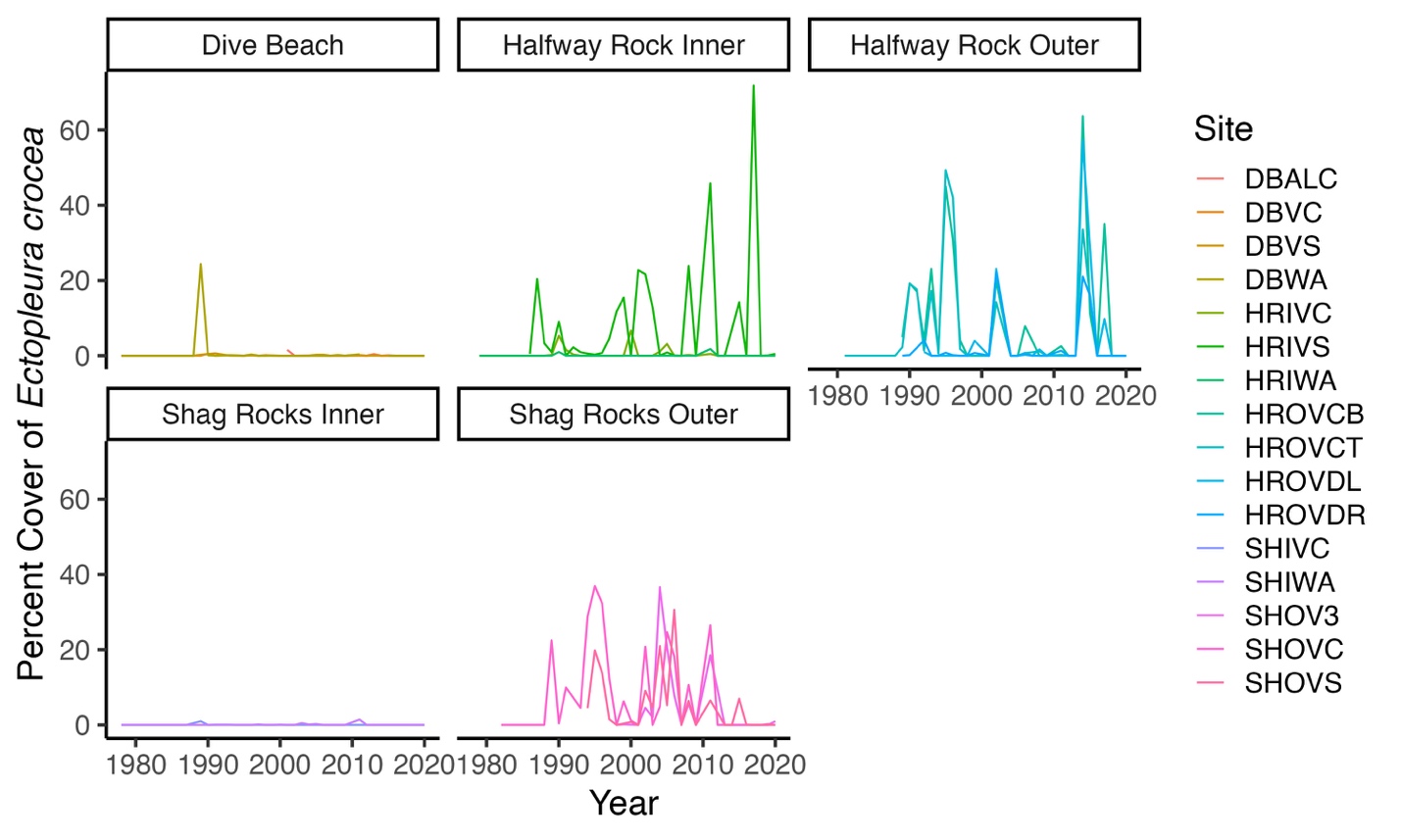

N.

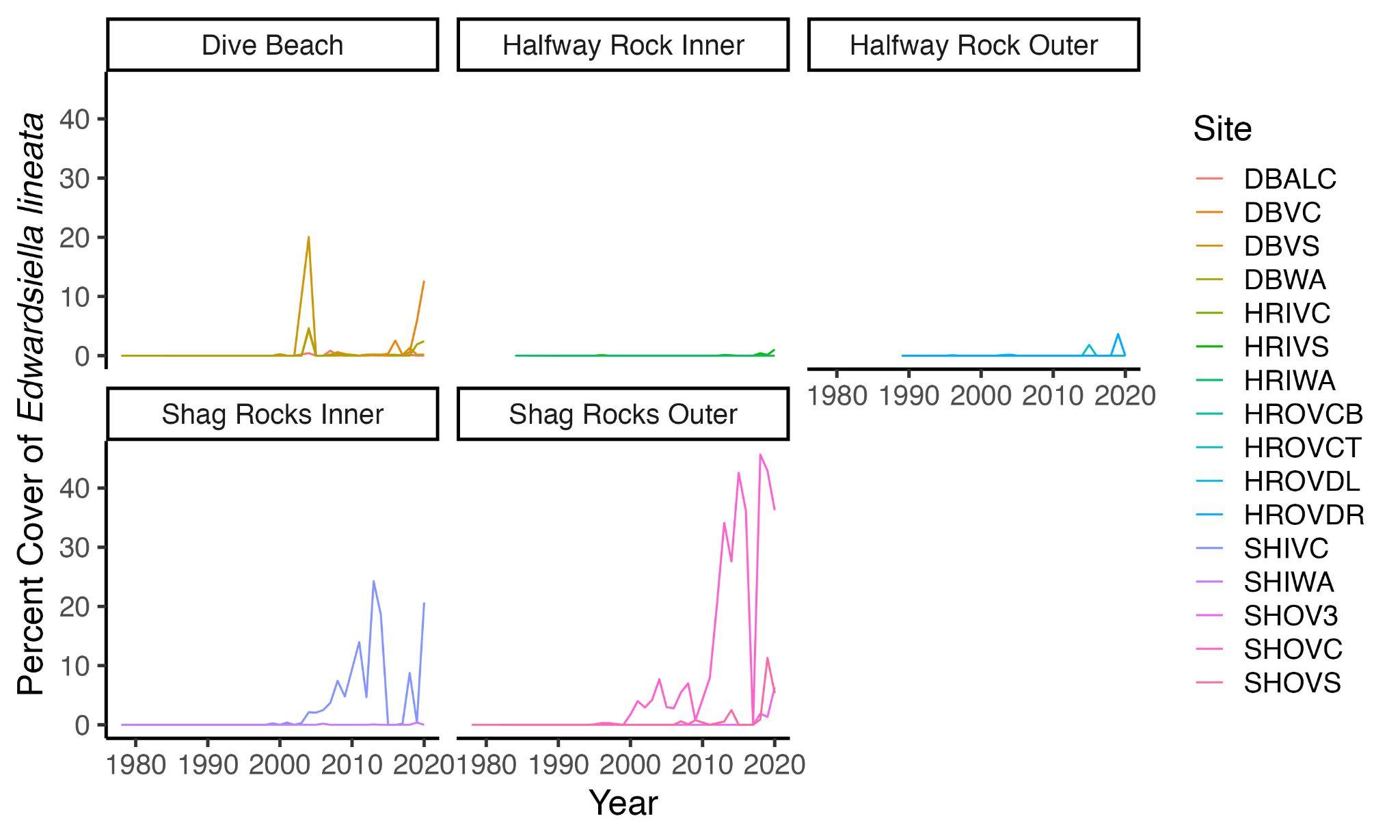

O.

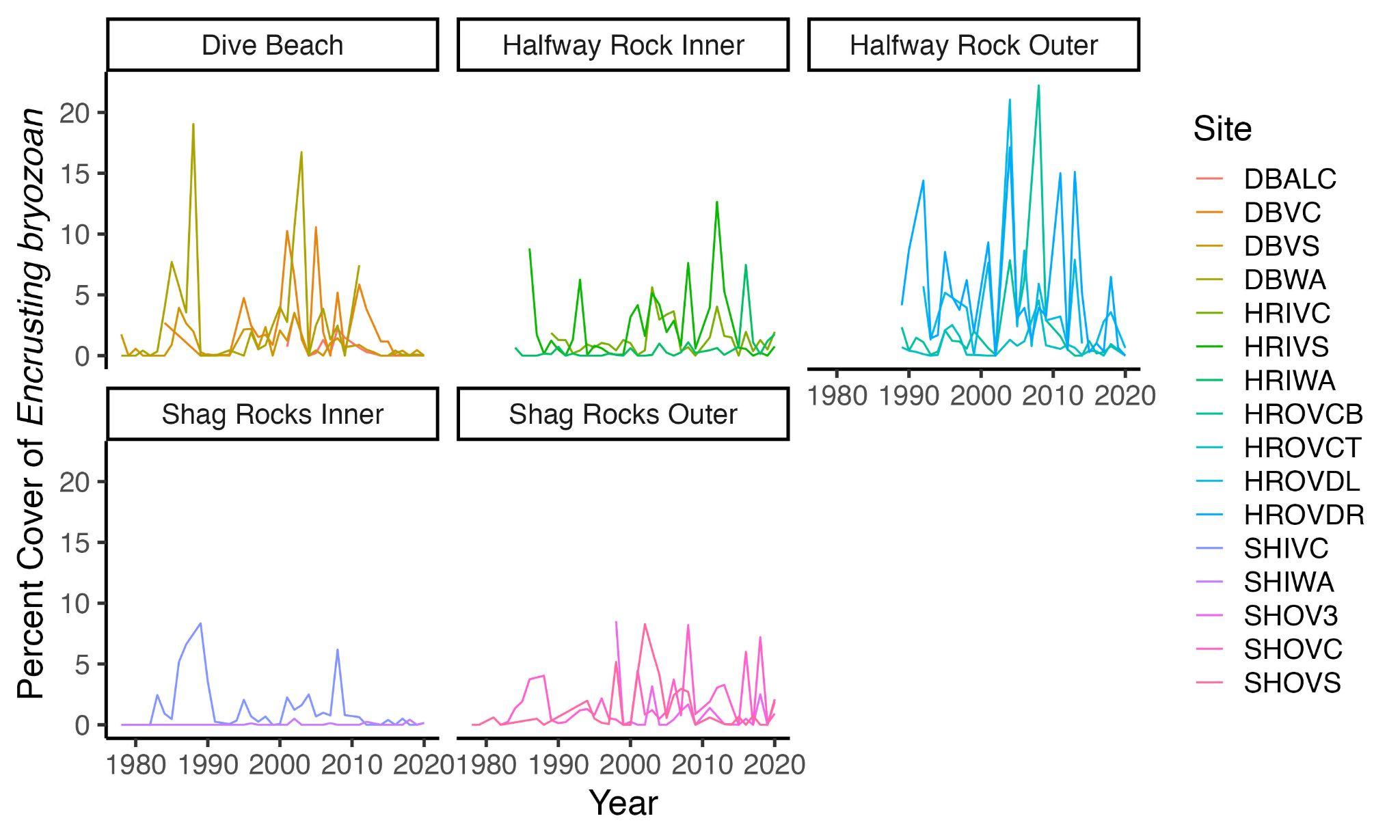

P.

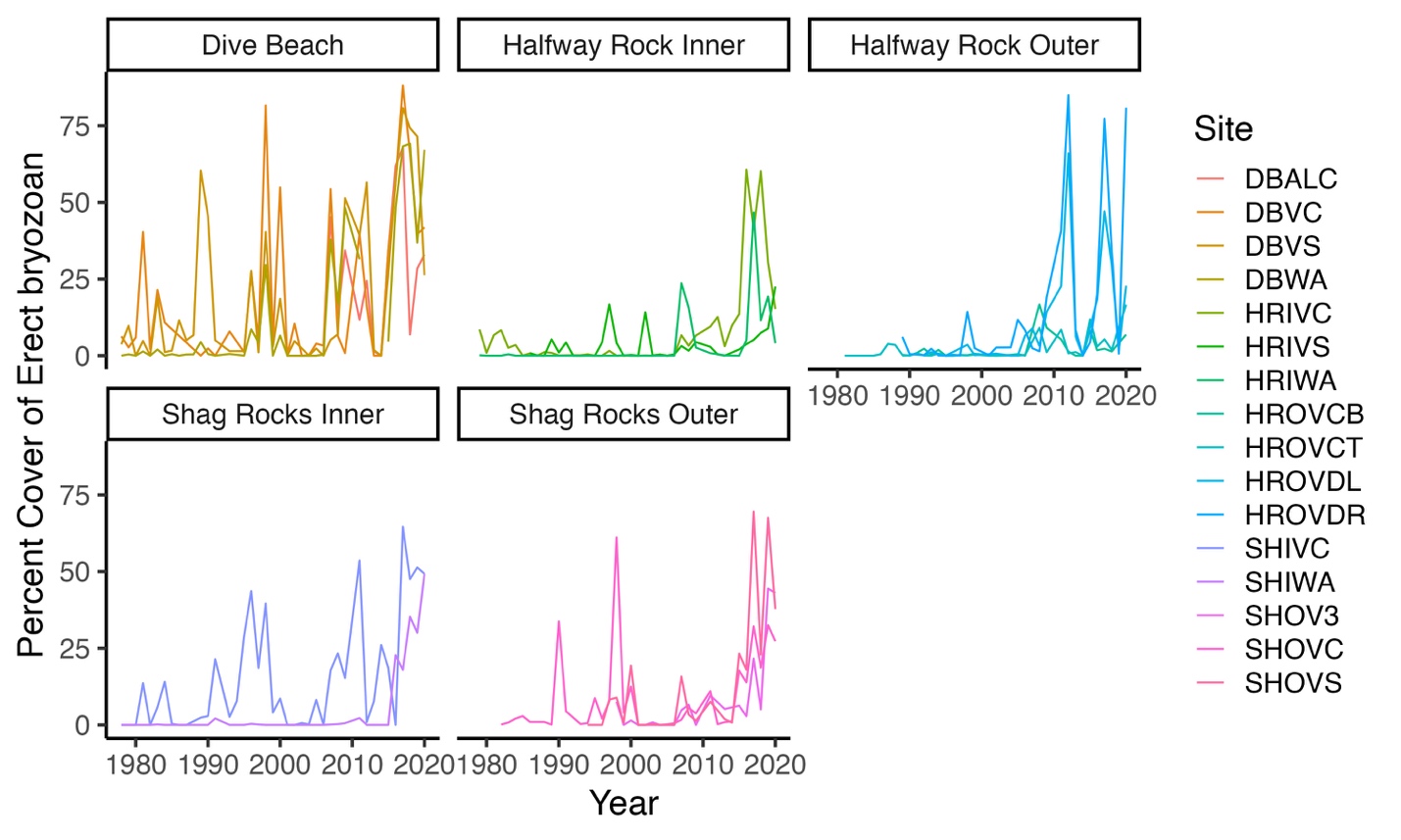

Q.

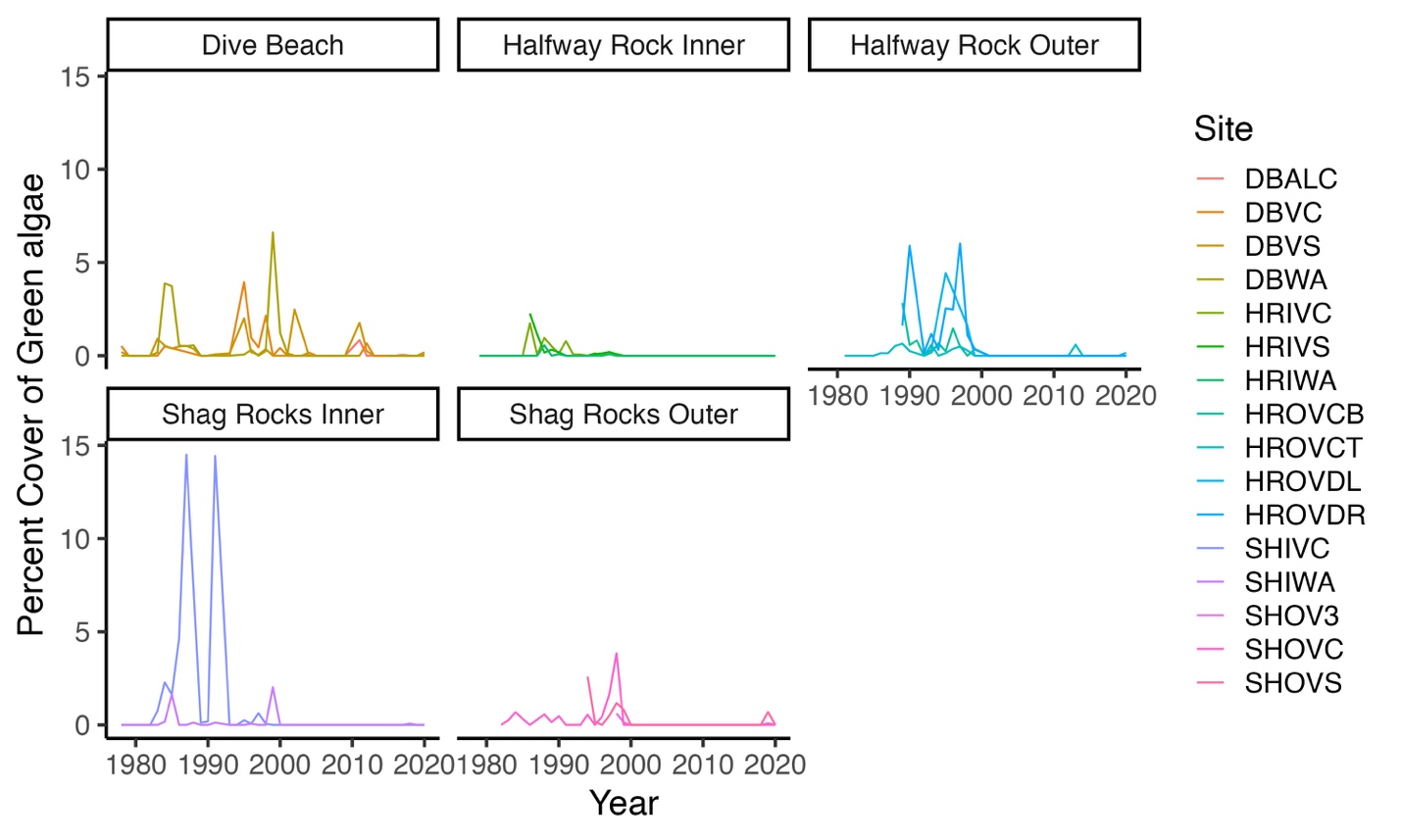

R.

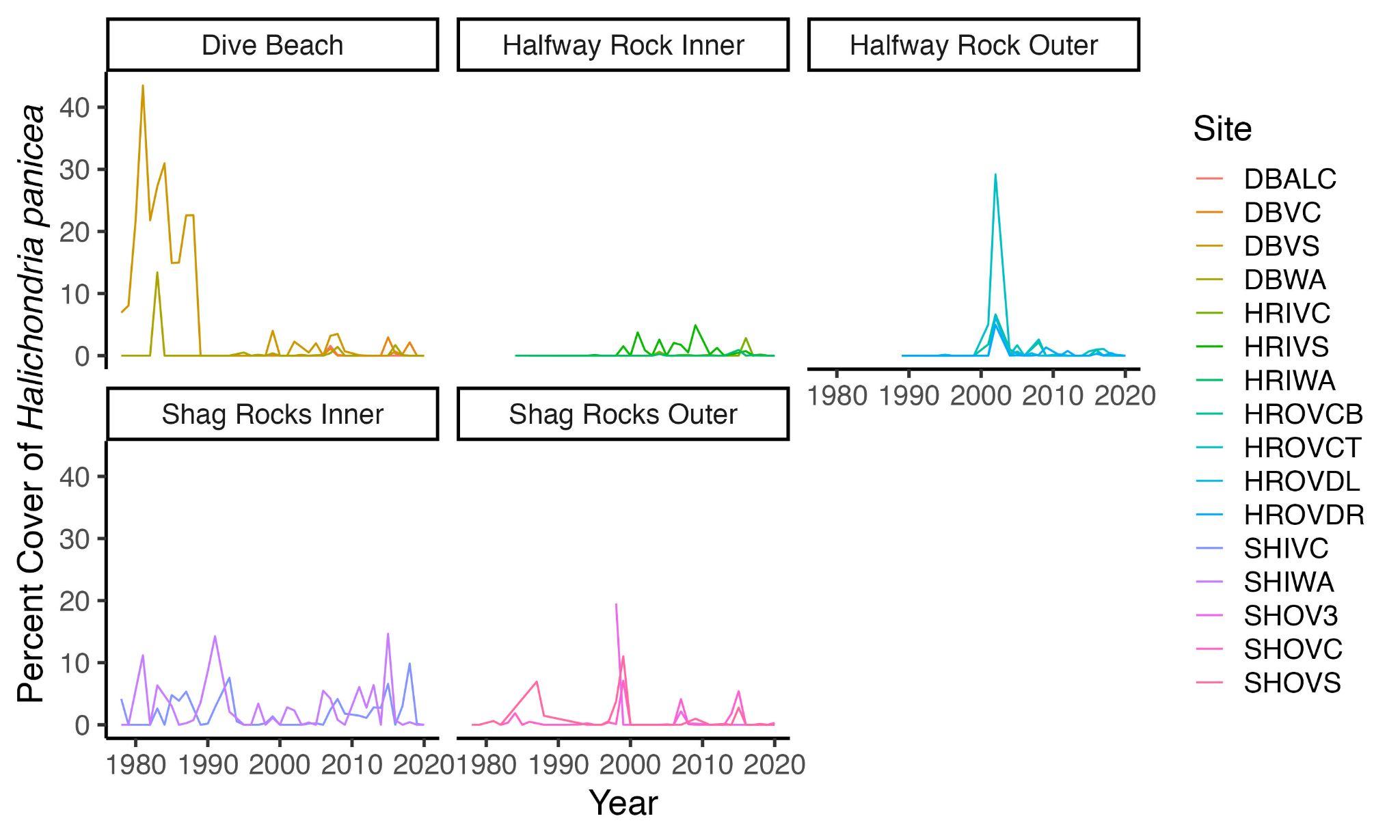

S.

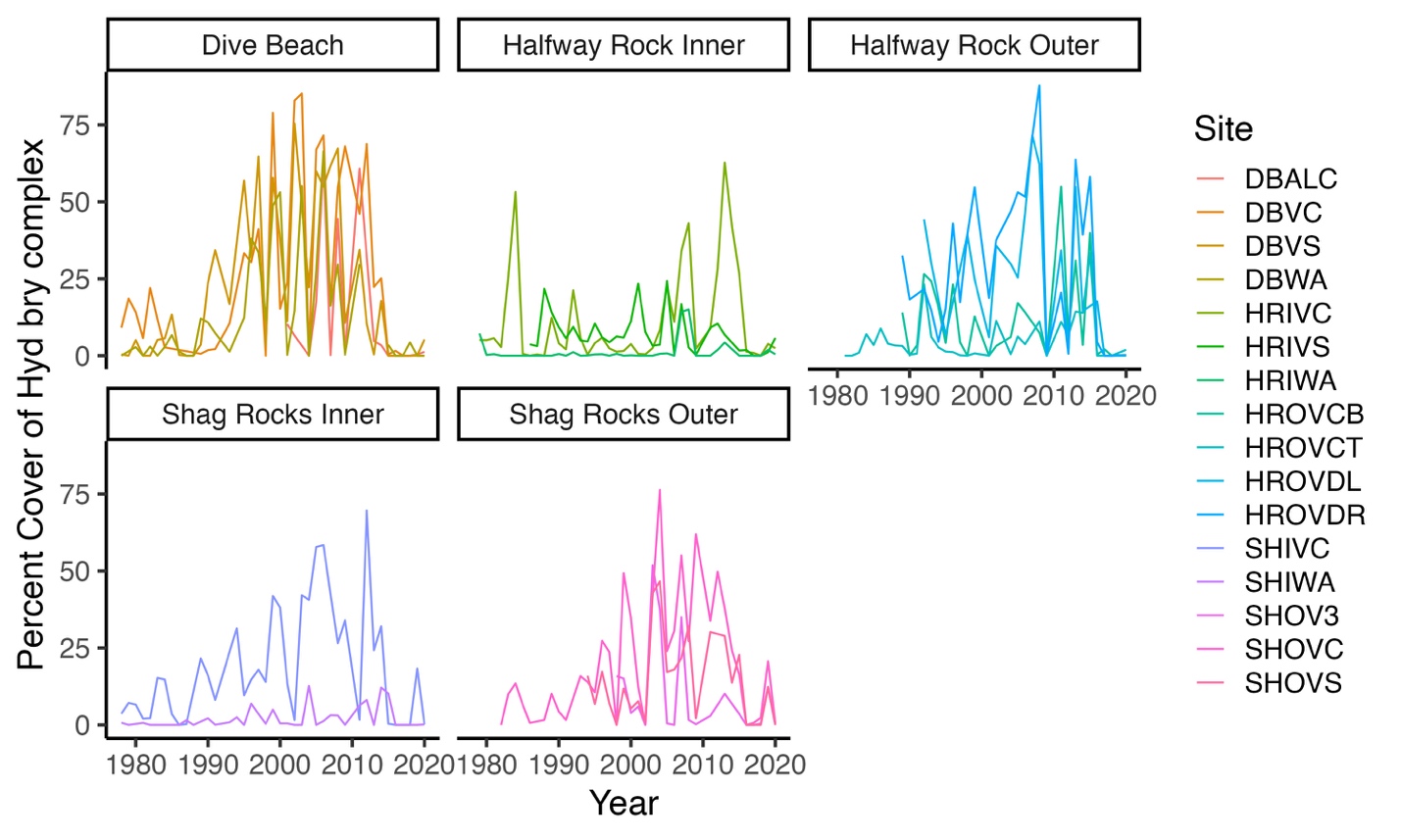

T.

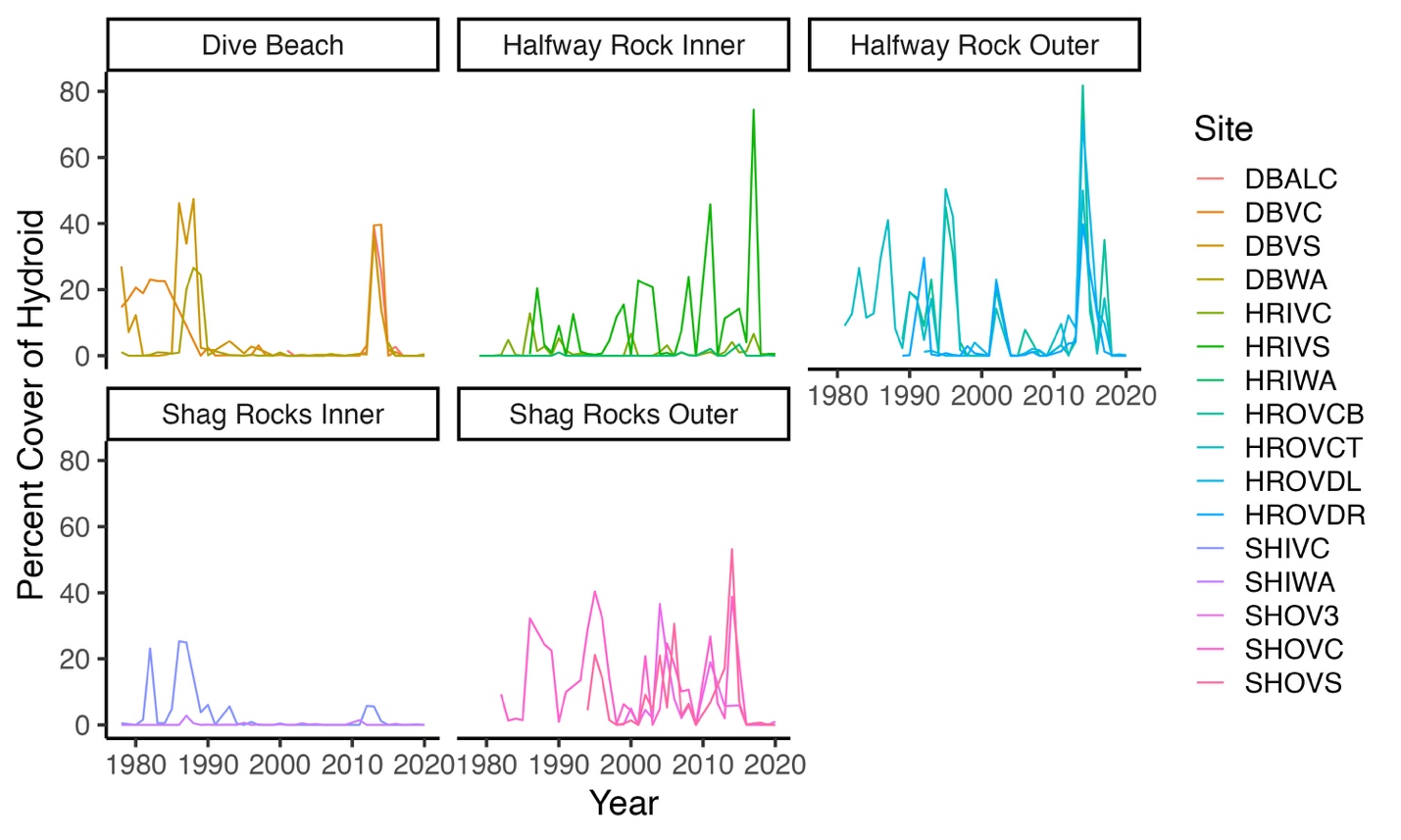

U.

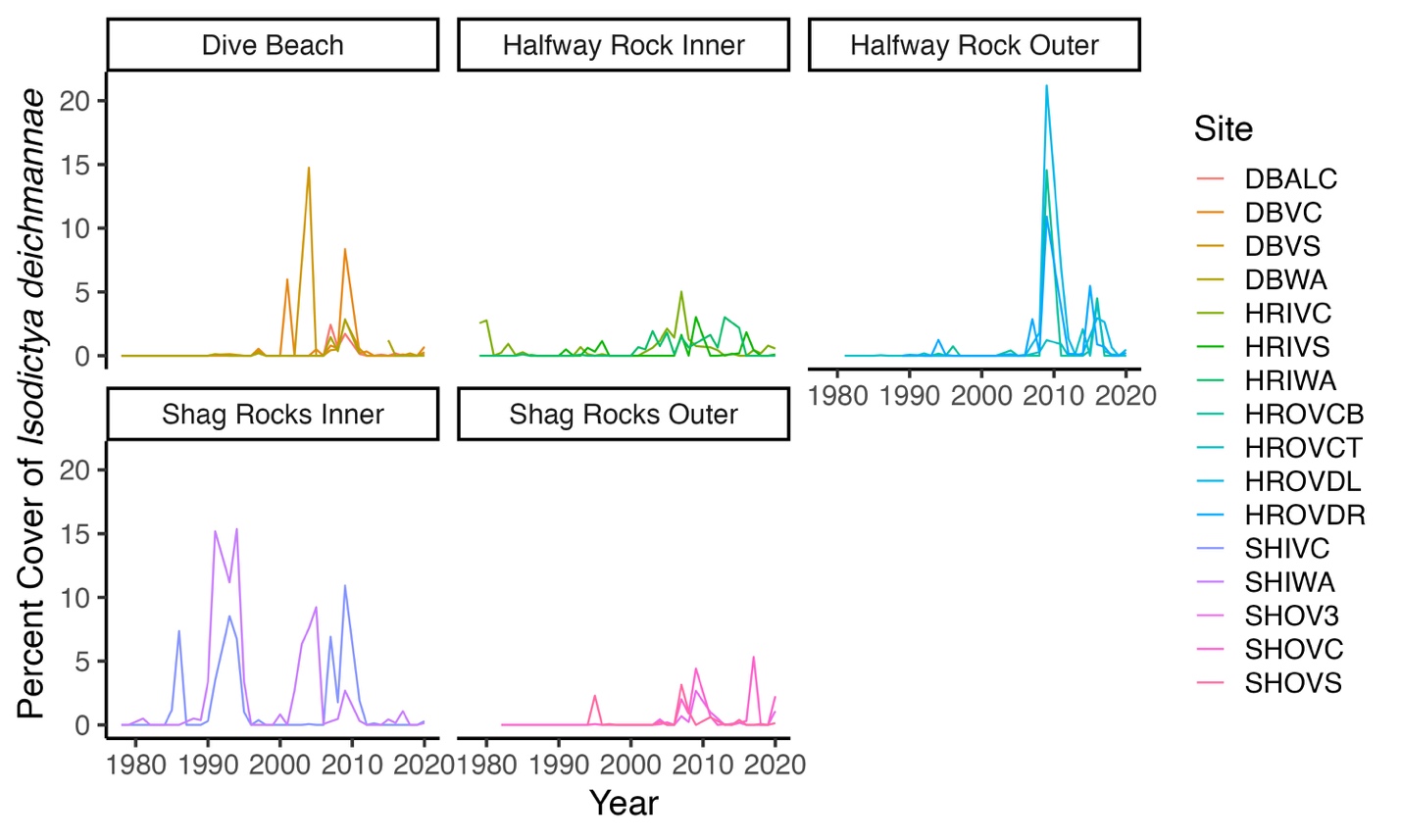

V.

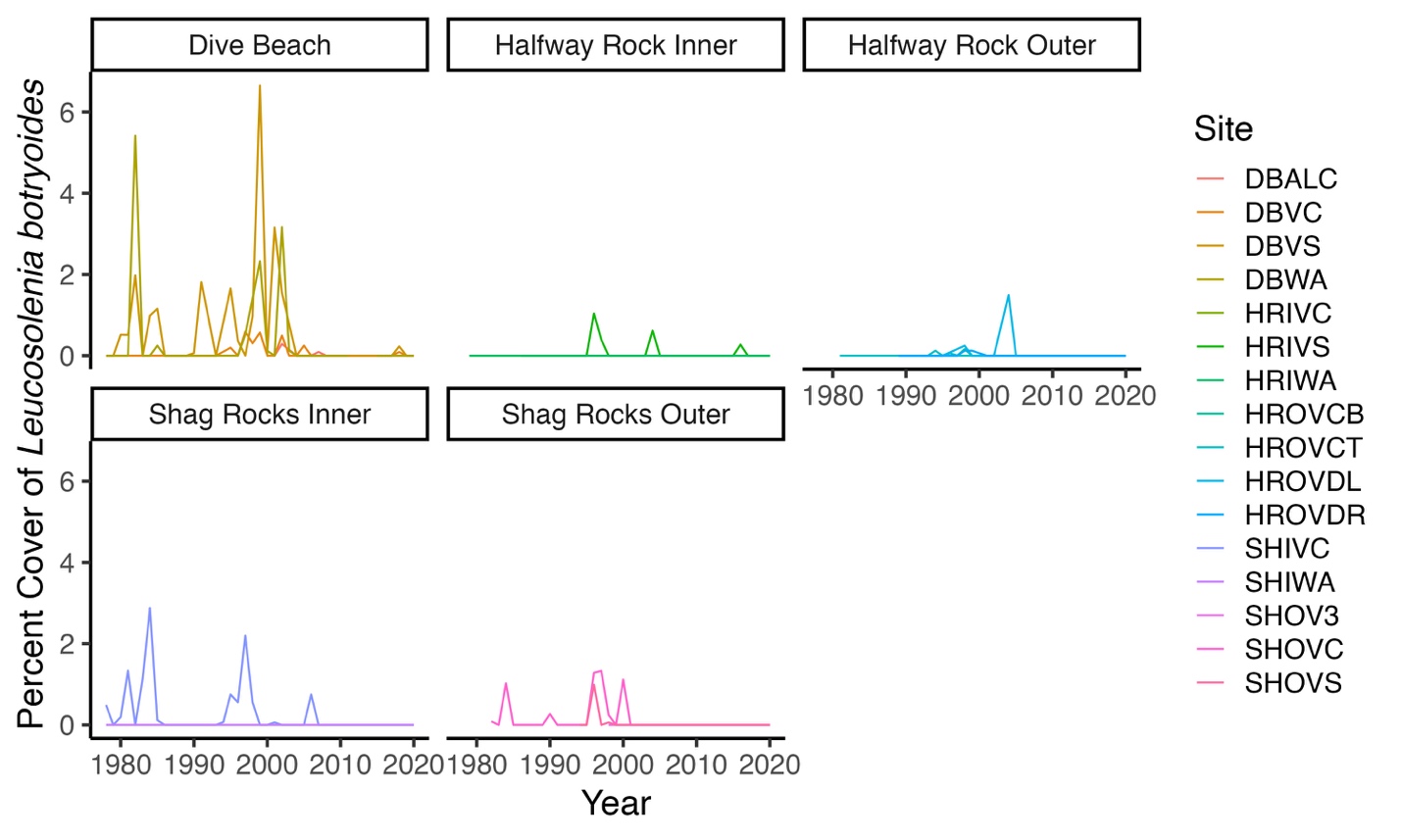

W.

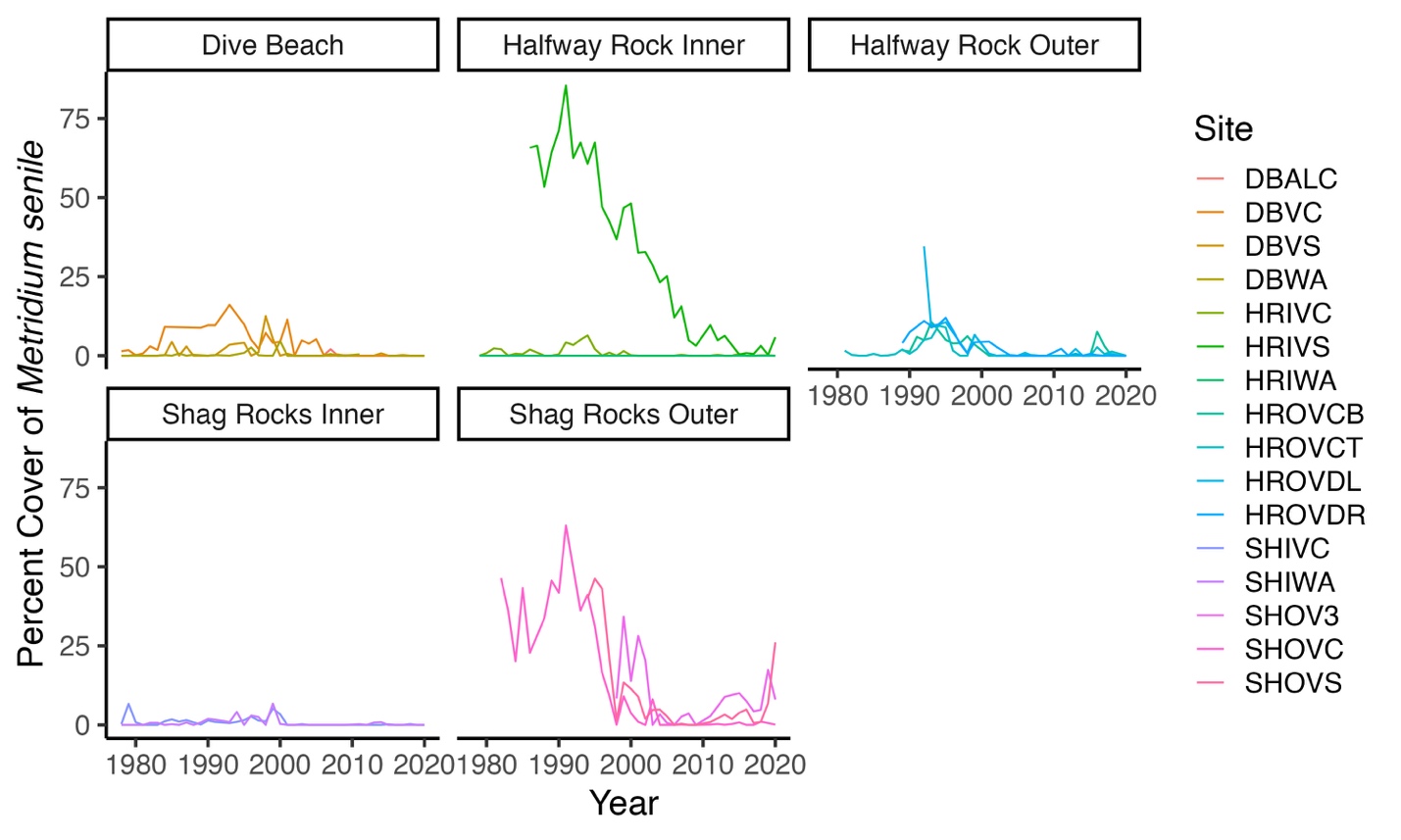

X.

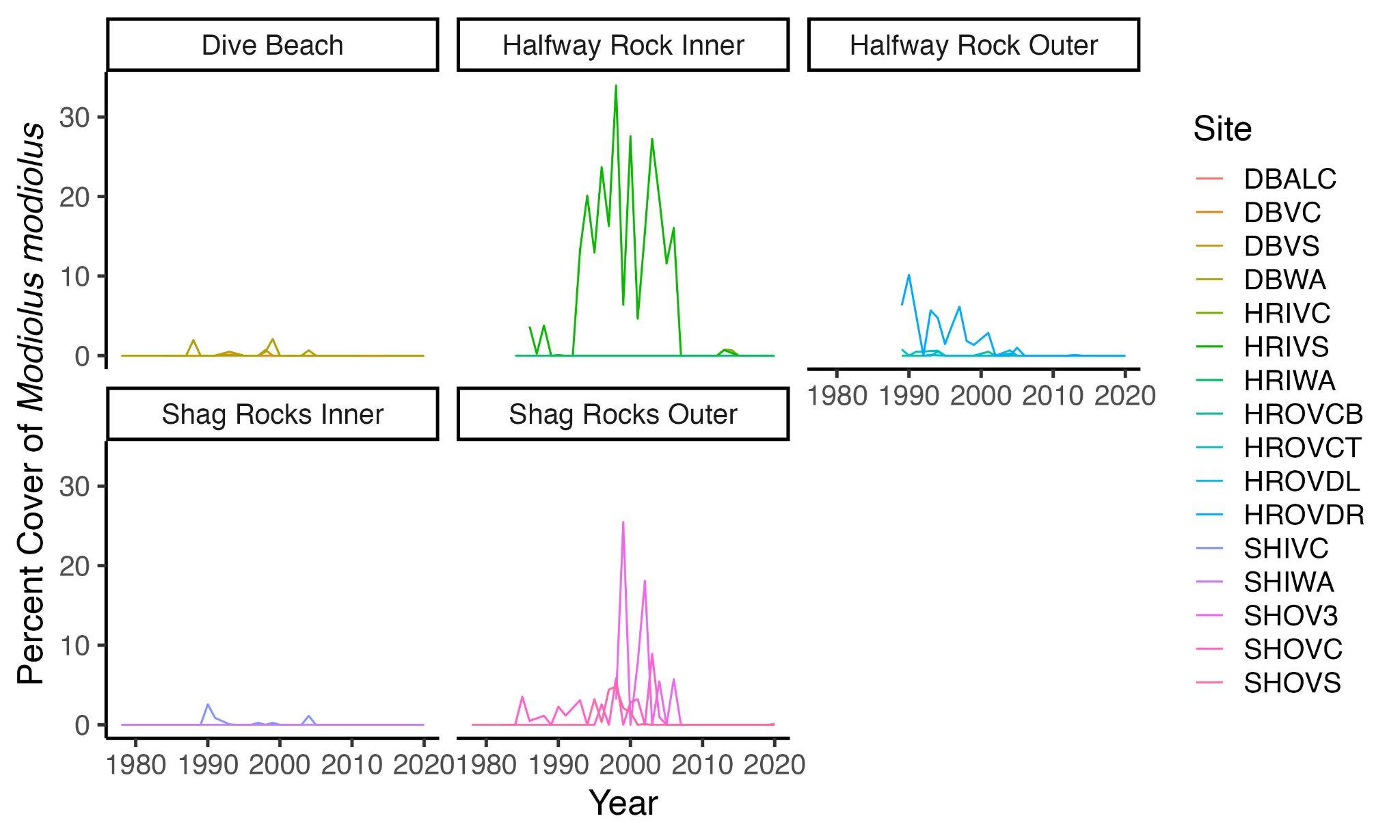

Y.

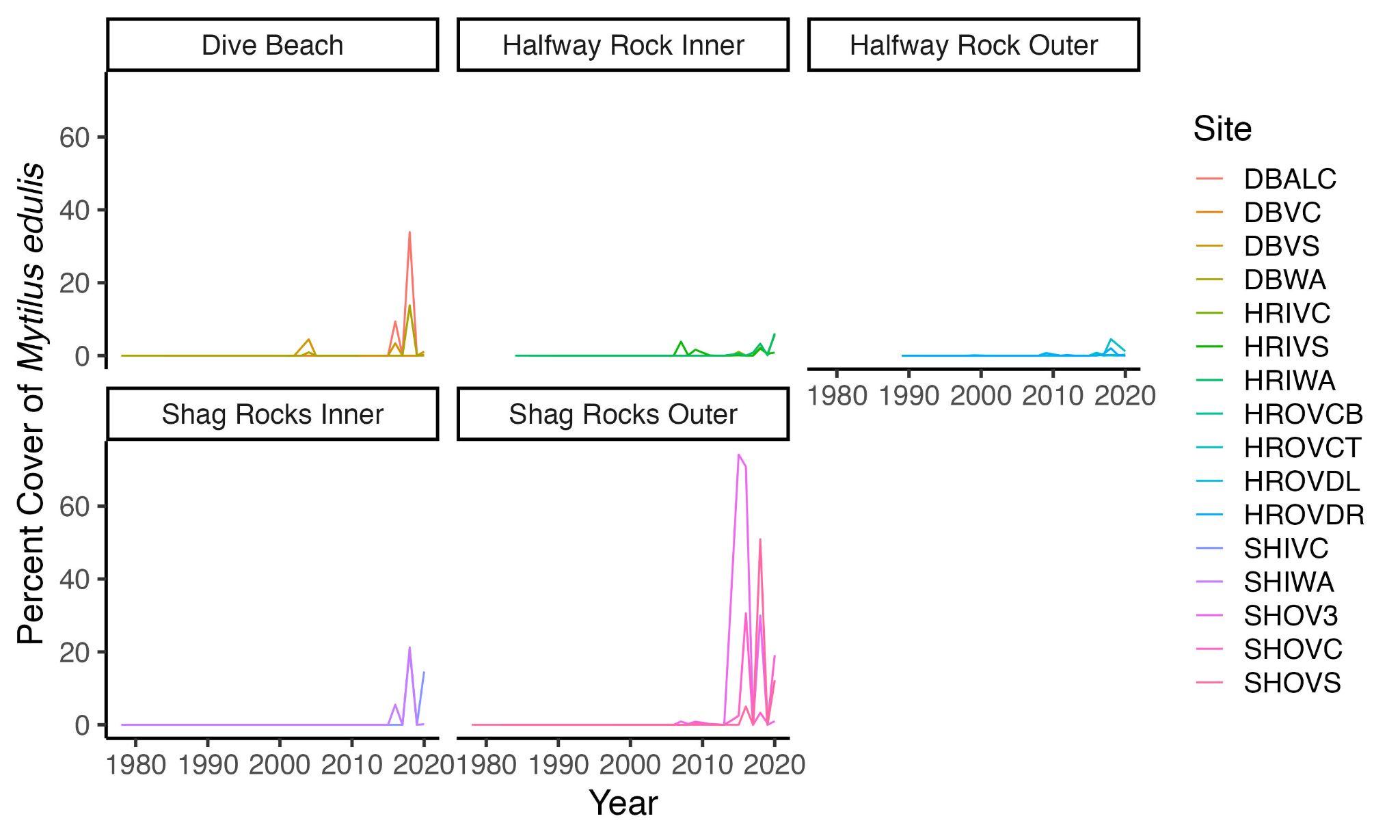

Z.

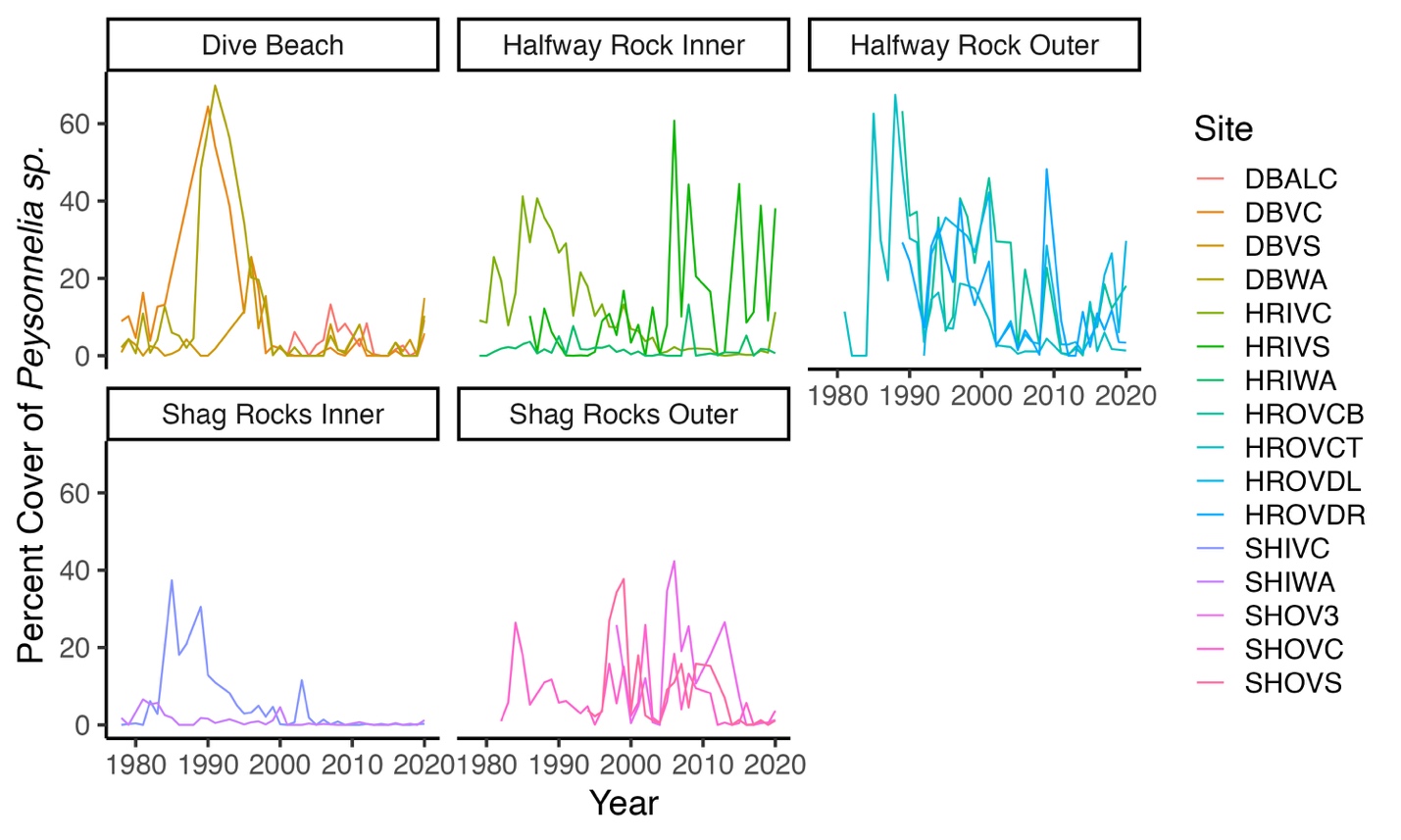

AA.

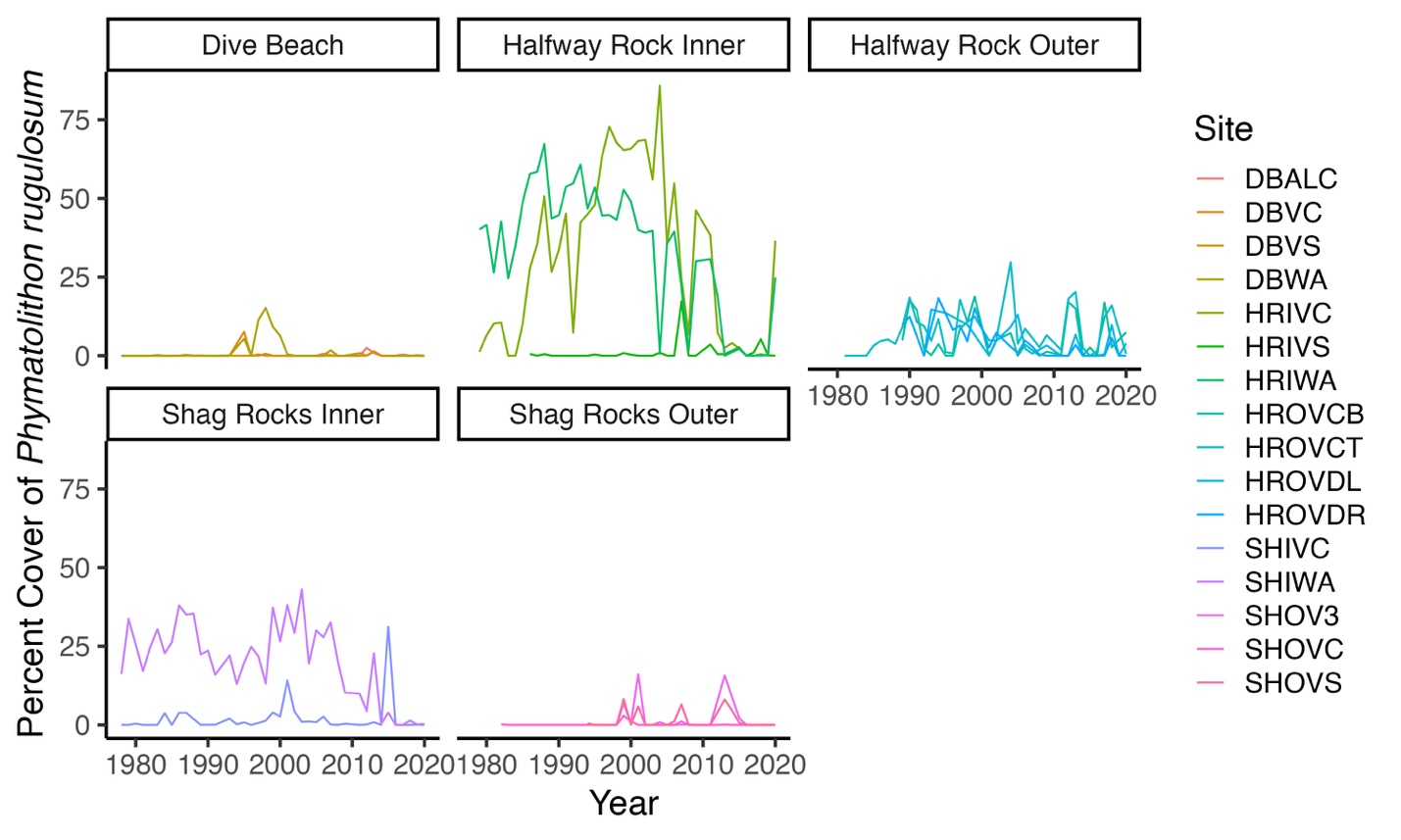

AB.

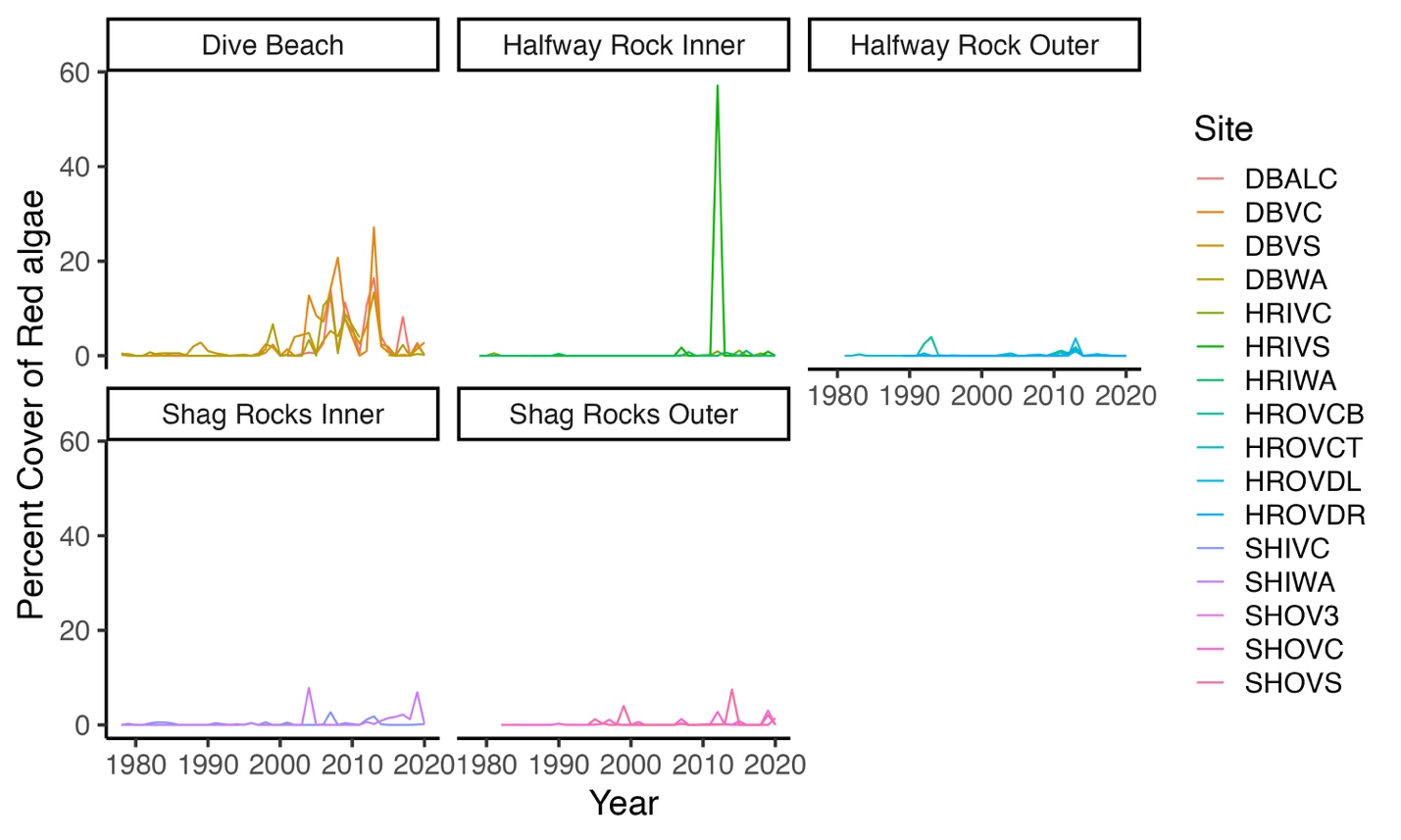

AC.

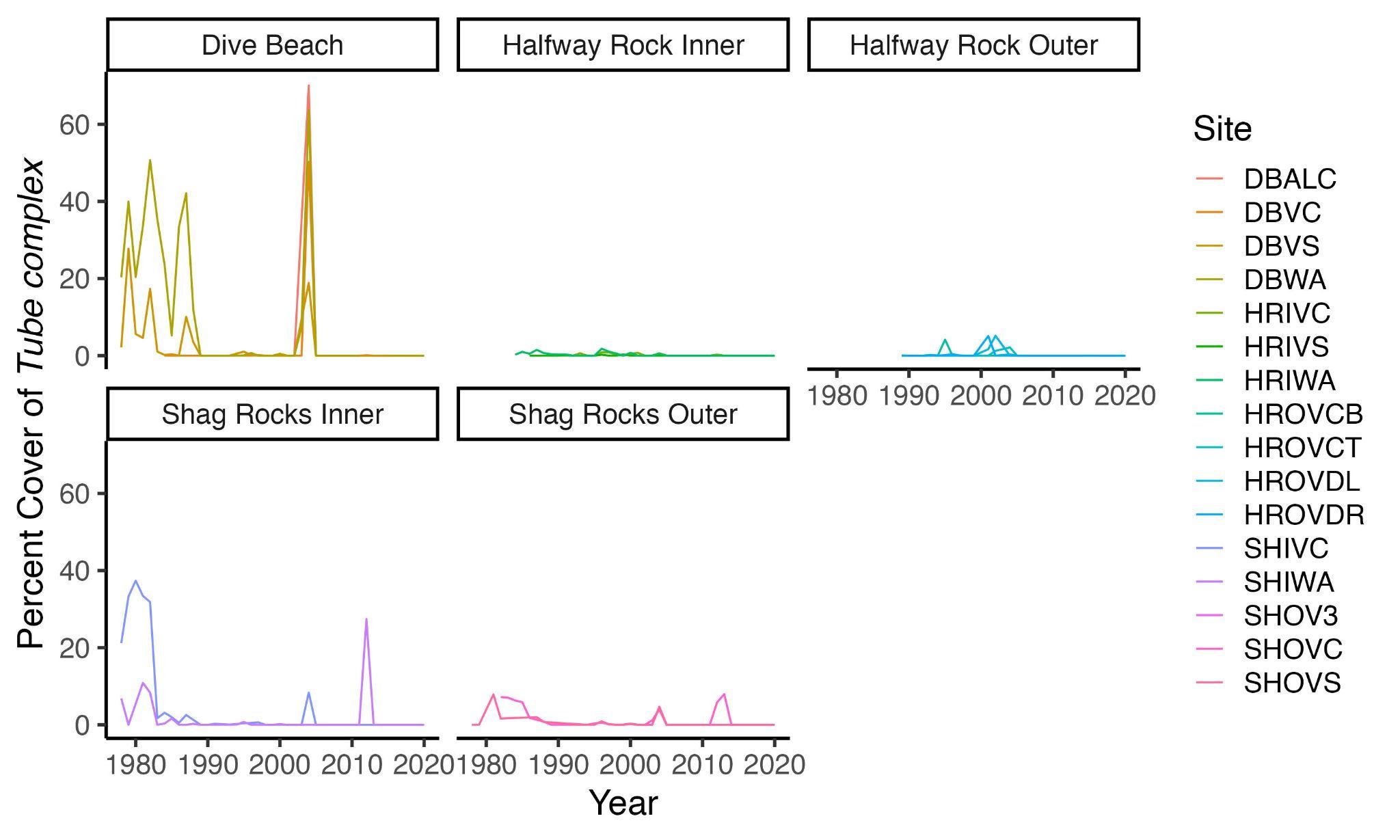

Table 3. Phyla, latitude, thermal maxima, and Bayesian statistics for all species and combined species used in this study. For latitudinal range, species with a southern range north of 40 degrees north are considered northern, those with a southern range south of 30 degrees are southern, and those with a southern range between 30-40 are intermediate (along east coast of North America).

| **Thermal Maxima Temperature Group** | **Rate of Change** | **Std. Error** | **Bayesian Credible Interval Low** | **Bayesian Credible Interval High** | **R2** | **Notes** |
| --- | --- | --- | --- | --- | --- | --- |
| Cold-affinity | 0.0684 | 0.0092 | 0.0505 | 0.0869 | 0.1376 |  |
| Cool-affinity | 0.0196 | 0.0036 | 0.0126 | 0.0266 | 0.2326 | *B. glaciale* was previously *Lithothamnion glaciale* |
| Cool-affinity | -0.0207 | 0.0041 | -0.0286 | -0.0127 | 0.2119 | *P. rugulosum* is a synonym of *P. scabriusculum* |
| Cool-affinity | -0.0101 | 0.0108 | -0.0308 | 0.0118 | 0.0877 |  |
| Cool-affinity | -0.0101 | 0.0058 | -0.0214 | 0.0015 | 0.0950 |  |
| Warm-affinity | 0.0101 | 0.0089 | -0.0068 | 0.0282 | 0.1284 |  |
| Cool-affinity | -0.0260 | 0.0087 | -0.0429 | -0.0085 | 0.1833 |  |
| Cool-affinity | -0.0440 | 0.0045 | -0.0528 | -0.0349 | 0.2367 |  |
| Cold-affinity | -0.0010 | 0.0080 | -0.0257 | 0.0052 | 0.1723 |  |
| Warm-affinity | 0.0584 | 0.0082 | 0.0431 | 0.0747 | 0.1309 | non-native |
| Cool-affinity | 0.0024 | 0.0065 | -0.0103 | 0.0151 | 0.0413 |  |
| Cool-affinity | 0.1975 | 0.0176 | 0.1643 | 0.2333 | 0.3635 | non-native |
| Warm-affinity | 0.0774 | 0.0119 | 0.0547 | 0.1018 | 0.1011 | non-native |
| Warm-affinity | 0.1155 | 0.0171 | 0.0832 | 0.1496 | 0.2521 |  |
| Cool-affinity | -0.0285 | 0.0090 | -0.0465 | -0.0118 | 0.1269 |  |
| Warm-affinity | 0.1693 | 0.0197 | 0.1318 | 0.2094 | 0.1594 |  |
| Cool-affinity | -0.0647 | 0.0070 | -0.0785 | -0.0511 | 0.2010 |  |
| Warm-affinity | -0.0074 | 0.0065 | -0.0201 | 0.0054 | 0.0949 |  |
| Warm-affinity | 0.1488 | 0.0107 | 0.1282 | 0.1697 | 0.1880 |  |
| Cool-affinity | -0.0653 | 0.0052 | -0.0753 | -0.0550 | 0.2303 |  |
| Cool-affinity | -0.0053 | 0.0045 | -0.0143 | 0.0035 | 0.0413 |  |
| NA | 0.0486 | 0.0044 | 0.0401 | 0.0573 | 0.2065 | *Bugula turrita, B. simples*, and others |
| NA | 0.0000 | 0.0038 | -0.0073 | 0.0075 | 0.0792 | *Shizomavella spp*., and others |
| NA | -0.0509 | 0.0073 | -0.0657 | -0.0368 | 0.1151 | mats of small filamentous algae |
| NA | -0.0094 | 0.0047 | -0.0186 | -0.0001 | 0.0338 | *Obelia spp., Ectopleura spp*. and others |
| NA | 0.0001 | 0.0041 | -0.0078 | 0.0080 | 0.0617 | seidment bound by stalks of hydroids and bryozoans (brown fuzz) |
| NA | -0.0306 | 0.0039 | -0.0383 | -0.0230 | 0.1453 | *Peysonnelia sp*. Is now *Waernia mirabilis* R.T.Wilce, Maggs & Sears, 2003 |
| NA | 0.0264 | 0.0050 | 0.0165 | 0.0364 | 0.0803 | filamentous and foliose algae, often Chondrus crispus |
| NA | -0.0972 | 0.0086 | -0.1145 | -0.0810 | 0.2026 | seidment bound by polychaete and amphipod (*Jassa falcata*) tubes |

| **Species (possible species)** | **Phylum - group** | **Southern Limit** | **Northern Limit** | **Latitude Group** | **Thermal Maxima Temperature Group** | **Rate of Change** | **Std. Error** | **Bayesian Credible Interval Low** | **Bayesian Credible Interval High** | **R^2^** | **Notes** |
| --- | --- | --- | --- | --- | --- | --- | --- | --- | --- | --- | --- |
| **Encrusting Red Algae** |  |  |  |  |  |  |  |  |  |  |  |
| *Clathromorphum circumscriptum* (Strömfelt) Foslie, 1898 | Rhodophyta - coralline | 41.93 | 66.02 | Northern | Cool-affinity | 0.0684 | 0.0092 | 0.0505 | 0.0869 | 0.1376 |  |
| *Boreolithothamnion glaciale (*Kjellman) P.W. Gabrielson, Maneveldt, Hughey & V. Peña 2023 | Rhodophyta - coralline | 42.25 | 69.24 | Northern | Warm-affinity | 0.0196 | 0.0036 | 0.0126 | 0.0266 | 0.2326 | *B. glaciale* was previously *Lithothamnion glaciale* |
| *Phymatolithon* *rugulosum* W.H. Adey 1964 | Rhodophyta - coralline | 44.95 | 45.22 | Northern | Warm-affinity | -0.0207 | 0.0041 | -0.0286 | -0.0127 | 0.2119 | *P. rugulosum* is a synonym of *P. scabriusculum* |
| **Sponges** |  |  |  |  |  |  |  |  |  |  |  |
| *Cliona celata* Grant, 1826 | Porifera | 5.15 | 45.17 | Southern | Warm-affinity | -0.0101 | 0.0108 | -0.0308 | 0.0118 | 0.0877 |  |
| *Halichondria panicea* Pallas, 1766 | Porifera | 8.81 | 60.48 | Southern | Warm-affinity | -0.0101 | 0.0058 | -0.0214 | 0.0015 | 0.0950 |  |
| *Isodictya deichmannae* de Laubenfels, 1949 | Porifera | 40.41 | 46.03 | Northern | Hot-affinity | 0.0101 | 0.0089 | -0.0068 | 0.0282 | 0.1284 |  |
| *Leucosolenia botryoides* Ellis & Solander, 1786 | Porifera | 32.52 | 52.59 | Intermediate | Warm-affinity | -0.0260 | 0.0087 | -0.0429 | -0.0085 | 0.1833 |  |
| **Tunicates** |  |  |  |  |  |  |  |  |  |  |  |
| *Aplidium glabrum* Verrill, 1871 | Chordata - ascidian | 32.52 | 63.67 | Intermediate | Warm-affinity | -0.0440 | 0.0045 | -0.0528 | -0.0349 | 0.2367 |  |
| *Boltenia echinata* Linnaeus, 1767 | Chordata - ascidian | 42.01 | 74.68 | Northern | Cool-affinity | -0.0010 | 0.0080 | -0.0257 | 0.0052 | 0.1723 |  |
| *Botrylloides violaceus* Oka, 1927 | Chordata - ascidian | 10.69 | 46.92 | Southern | Hot-affinity | 0.0584 | 0.0082 | 0.0431 | 0.0747 | 0.1309 | non-native |
| *Didemnum albidum* Verrill, 1871 | Chordata - ascidian | 38.65 | 65.95 | Intermediate | Warm-affinity | 0.0024 | 0.0065 | -0.0103 | 0.0151 | 0.0413 |  |
| *Didemnum vexillum* Kott, 2002 | Chordata - ascidian | 40.1 | 44.98 | Northern | Warm-affinity | 0.1975 | 0.0176 | 0.1643 | 0.2333 | 0.3635 | non-native |
| *Diplosoma listerianum* Milne Edwards, 1841 | Chordata - ascidian | 11 | 51.85 | Southern | Hot-affinity | 0.0774 | 0.0119 | 0.0547 | 0.1018 | 0.1011 | non-native |
| **Molluscs** |  |  |  |  |  |  |  |  |  |  |  |
| *Anomia simplex* d'Orbigny, 1853 | Mollusca | 4.45 | 46.69 | Southern | Hot-affinity | 0.1155 | 0.0171 | 0.0832 | 0.1496 | 0.2521 |  |
| *Modiolus modiolus* Linnaeus, 1758 | Mollusca | 4.17 | 61.57 | Southern | Warm-affinity | -0.0285 | 0.0090 | -0.0465 | -0.0118 | 0.1269 |  |
| *Mytilus edulis* Linnaeus, 1758 | Mollusca | 10.08 | 82.19 | Southern | Hot-affinity | 0.1693 | 0.0197 | 0.1318 | 0.2094 | 0.1594 |  |
| **Cnidarians** |  |  |  |  |  |  |  |  |  |  |  |
| *Alcyonium siderium* Verrill, 1922 | Cnidaria - octocoral | 42.34 | 44.91 | Northern | Warm-affinity | -0.0647 | 0.0070 | -0.0785 | -0.0511 | 0.2010 |  |
| *Ectopleura crocea* Agassiz, 1862 | Cnidaria - hydrozoan | 29.67 | 44.99 | Southern | Hot-affinity | -0.0074 | 0.0065 | -0.0201 | 0.0054 | 0.0949 |  |
| *Edwardsiella lineata* Verrill, 1873 | Cnidaria - anemone | 37.2 | 43.13 | Intermediate | Hot-affinity | 0.1488 | 0.0107 | 0.1282 | 0.1697 | 0.1880 |  |
| *Metridium senile* Linneaus, 1761 | Cnidaria - anemone | 16.4 | 68.71 | Southern | Warm-affinity | -0.0653 | 0.0052 | -0.0753 | -0.0550 | 0.2303 |  |
| **Other** |  |  |  |  |  |  |  |  |  |  |  |
| *Balanus balanus* Linnaeus, 1758 | Arthropoda - crustacean | 22.05 | 76.85 | Southern | Warm-affinity | -0.0053 | 0.0045 | -0.0143 | 0.0035 | 0.0413 |  |
| **Combined Groups** |  |  |  |  |  |  |  |  |  |  |  |
| Erect Bryozoans | Bryozoa/Ectoprocta - bryozoans | NA | NA | NA | NA | 0.0486 | 0.0044 | 0.0401 | 0.0573 | 0.2065 | Bugula turrita, B. simples, and others |
| Encrusting Bryozoans | Bryozoa/Ectoprocta - bryozoans | NA | NA | NA | NA | 0.0000 | 0.0038 | -0.0073 | 0.0075 | 0.0792 | Shizomavella spp., and others |
| Green Algae | Chlorophyta | NA | NA | NA | NA | -0.0509 | 0.0073 | -0.0657 | -0.0368 | 0.1151 | mats of small filamentous algae |
| Hydroid | Cnidaria - hydrozoans | NA | NA | NA | NA | -0.0094 | 0.0047 | -0.0186 | -0.0001 | 0.0338 | Obelia spp., Ectopleura spp. and others |
| Hydroid/Bryozoan Complex | mixed | NA | NA | NA | NA | 0.0001 | 0.0041 | -0.0078 | 0.0080 | 0.0617 | seidment bound by stalks of hydroids and bryozoans (brown fuzz) |
| *Peysonnelia* sp. | Rhodophyrta - red crust | NA | NA | NA | NA | -0.0306 | 0.0039 | -0.0383 | -0.0230 | 0.1453 | *Peysonnelia sp*. Is now *Waernia mirabilis* R.T.Wilce, Maggs & Sears, 2003 |
| Red Algae | Rhodophyta | NA | NA | NA | NA | 0.0264 | 0.0050 | 0.0165 | 0.0364 | 0.0803 | filamentous and foliose algae, often Chondrus crispus |
| Tube Complex | mixed | NA | NA | NA | NA | -0.0972 | 0.0086 | -0.1145 | -0.0810 | 0.2026 | seidment bound by polychaete and amphipod (Jassa falcata) tubes |
